# Diverse Perceptual Representations Across Visual Pathways Emerge from A Single Objective

**DOI:** 10.1101/2025.07.22.664908

**Authors:** Yingtian Tang, Abdulkadir Gokce, Khaled Jedoui Al-Karkari, Daniel Yamins, Martin Schrimpf

## Abstract

How the human brain supports diverse behaviours has been debated for decades. The canonical account posits distinct pathways in perceptual processing, such as the “what” and “where/how” visual streams, though their developmental origins and interdependency remain contested. Here, we show that families of deep neural network models develop features that accurately predict hours of human neural and behavioural recordings. By using these models as proxies for the brain and systematically probing developmental objectives, we identify two fundamental computations: object and appearance-free motion recognition, which drive the visual processing hierarchy and explain away alternative accounts. Strikingly, a *single* objective underlies both: they emerge from optimising for understanding world dynamics, with their organisation highly distributed and continuous across cortex rather than segregating into stream-like modes. Our results suggest that the human brain’s ability to integrate complex perceptual information across seemingly distinct pathways may originate from the single goal of modelling the world.

## Introduction

In order to operate in an ever-changing environment, humans parse dynamic sensory information. From interpreting the hunt of a predator or the escape route of prey, to navigating cars and pedestrians in modern city environments, the brain continuously integrates a wealth of visual sensory input. Visual information is processed by the visual cortex [140, 54, 49, 103, 176, 165], where functionally specialised regions emerge for basic visual features [72, 19] as well as for complex patterns such as objects [32], faces [63, 162, 77], scenes [35, 35], and motion [6, 14, 160]. The concept of visual streams describes the computational hierarchy among these regions, with the classical account proposing a ventral (“what”) and a dorsal (“where/how”) stream based on neuroanatomical and lesion studies [119, 61, 29, 37, 118, 117, 43, 115]. Increasing evidence [114, 86, 48, 47, 141, 164, 85, 60, 152, 145] indicates that regions across both streams interact dynamically, with boundaries that are graded and continuous rather than sharply discrete. Efforts toward more detailed characterisation have yielded proposals of additional streams and finer subdivisions within streams [171, 140, 85, 142, 111, 131, 76, 87, 130], yet these accounts remain largely descriptive in nature. Despite outlining a qualitative computational hierarchy, these frameworks lack testable stimulus–response predictions and a unified quantitative explanation of the mechanisms and developmental origins underlying visual perception.

Advances in AI have led to artificial neural networks (ANNs) that achieve human-level performance, with internal machinery abstractly resembling biological neurons. The recent use of task-optimised AI models in systems neuroscience has revolutionised our ability to explain neural representations in distinct regions throughout the brain [174, 83, 147, 81, 126, 123, 84, 4, 148, 146, 149, 22, 59, 23, 3]. These models are optimised for computational tasks without fitting to brain data, and yet exhibit internal representations that are remarkably brain-like [173, 148, 146]. Linking task performance to neural alignment reveals evolutionary and developmental objectives that might have shaped brain representations: this task-driven modelling approach has established object categorisation as a key driver of static vision across the primate visual ventral stream [174, 147], with comparable findings in other systems, such as auditory [81, 95] and language processing [146, 22]. However, a crucial limitation has been the predominant focus on individual pathways and their corresponding objectives, with no coherent theory of the objectives that govern computations across the entire visual cortex and span multiple processing streams.

To address this gap, we here establish a normative computational framework of how dynamic vision is processed across visual cortex in the service of different task demands. We assess the internal representations in 92 pretrained deep neural networks and evaluate their alignment to hours of brain recordings across diverse datasets featuring dynamic stimuli, encompassing social interactions, biological motions, physical object dynamics, narrative sequences, and other facets of real-world perception. In order to determine which objective(s) best explain the computational nature of brain-like representations, we introduce a novel *multi-task relevance* method (Fig. 1a) that identifies combinations of objectives uniquely explaining selected brain representations.

**Figure 1.**
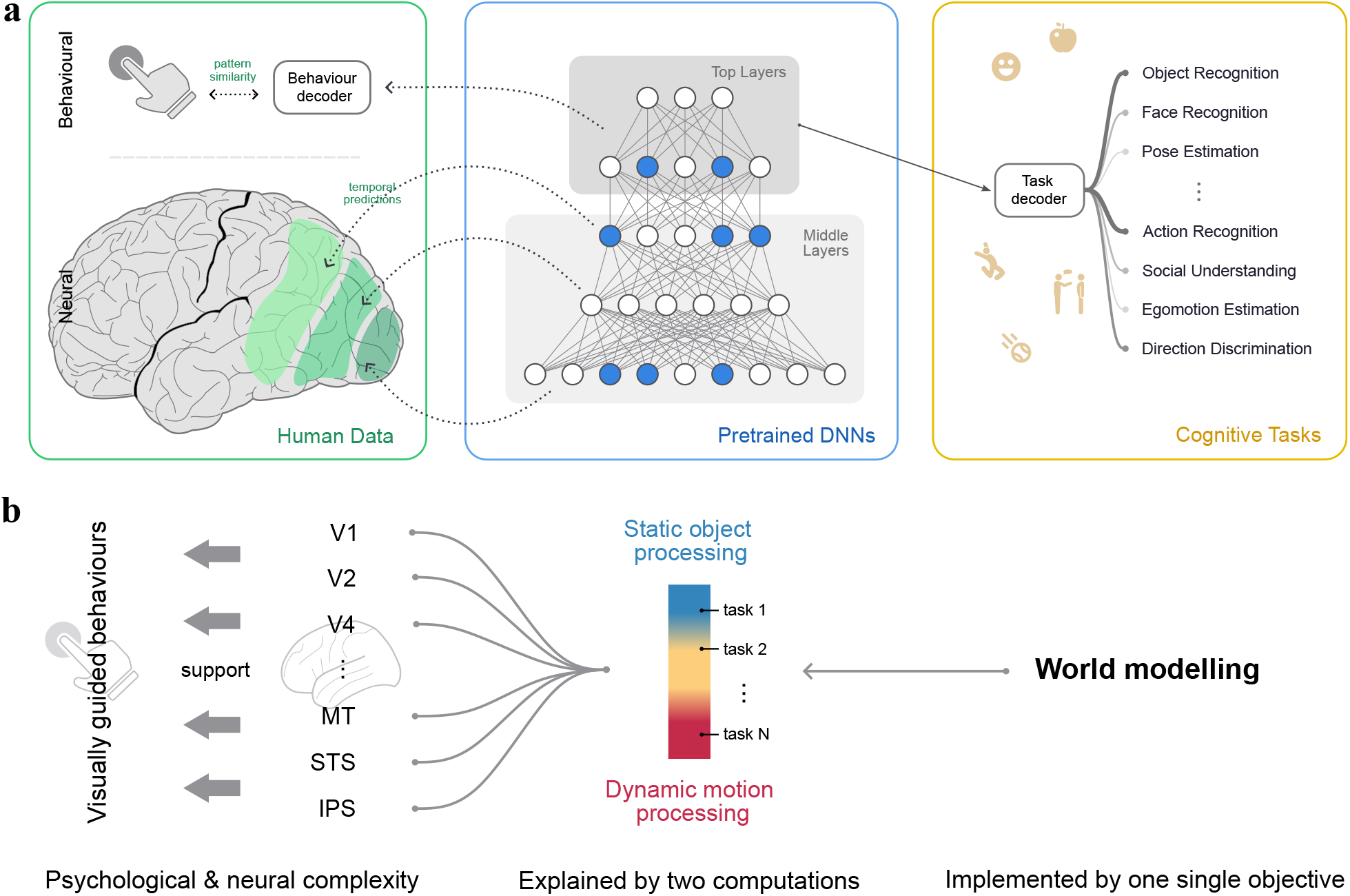
Multi-task computational framework for brain modelling. a. Models bridge neural, behavioural, and machine learning datasets: We test the alignment of representations from stimulus-computable models (*Middle*) to dynamic neural signals (fMRI) and behavioural recordings (*Left*), and connect brain alignment to task performance on diverse cognitive tasks (*Right* ; Fig. 3b), investigating hypotheses of which functional objectives shape representations in the brain. **b. Core claim:** We find that the psychological and neural complexity during perception in diverse brain areas is explained by two core tasks, which can be further reduced to a single overarching objective. Representations across various brain regions supporting many visually guided behaviours arise from two core tasks—object recognition and appearance-free motion recognition—which themselves originate from a single ‘world modelling’ objective.

Across all model families, we identify dynamic models as state-of-the-art in alignment to visual cortex activity and consequent behavioural choices, surpassing both static and traditional models, and accurately predicting neural activity on a voxel-wise, second-by-second basis. These models are optimised via a single computational objective such as prediction across time and space, yet perform well on a multitude of downstream tasks, and develop brain-like hierarchical representations across ventral and dorsal streams – which were previously thought to be shaped by distinct task demands (Fig. 1b). Our framework reveals two core computations that jointly drive this system-wise alignment: static object recognition and dynamic motion recognition. These computations map onto the visual cortex in a continuous manner, without evidence for sharply discrete streams. We further link them to behaviour, characterise the full hierarchy from dynamic stimuli through neural processing to action recognition outcomes, and reveal the hybrid object–motion processing throughout. Overall, our integrative approach unifies diverse neural and behavioural datasets into a single computational framework, providing a mechanistic and predictive account of visual processing across the cortex, one that challenges and advances beyond traditional descriptive theories.

## Results

At the core of our analyses is the evaluation of alignment between artificial neural network models (ANNs) and brain recordings. We use 5 fMRI brain datasets of video viewing, totalling 35.7 subject hours and approximately 2 hours of unique visual input, as well as an additional behavioural dataset of action understanding. The candidate models comprise a repertoire of ANNs pretrained on a diverse set of objectives (Methods B.2), including state-of-the-art deep learning approaches spanning static image classification to dynamic representation learning, such as masked video modelling. We assess model-to-brain alignment using a regression-based metric *across time*, which applies a time-invariant linear mapping from model units to brain voxels (Fig. 2a; Methods B.3).

**Figure 2.**
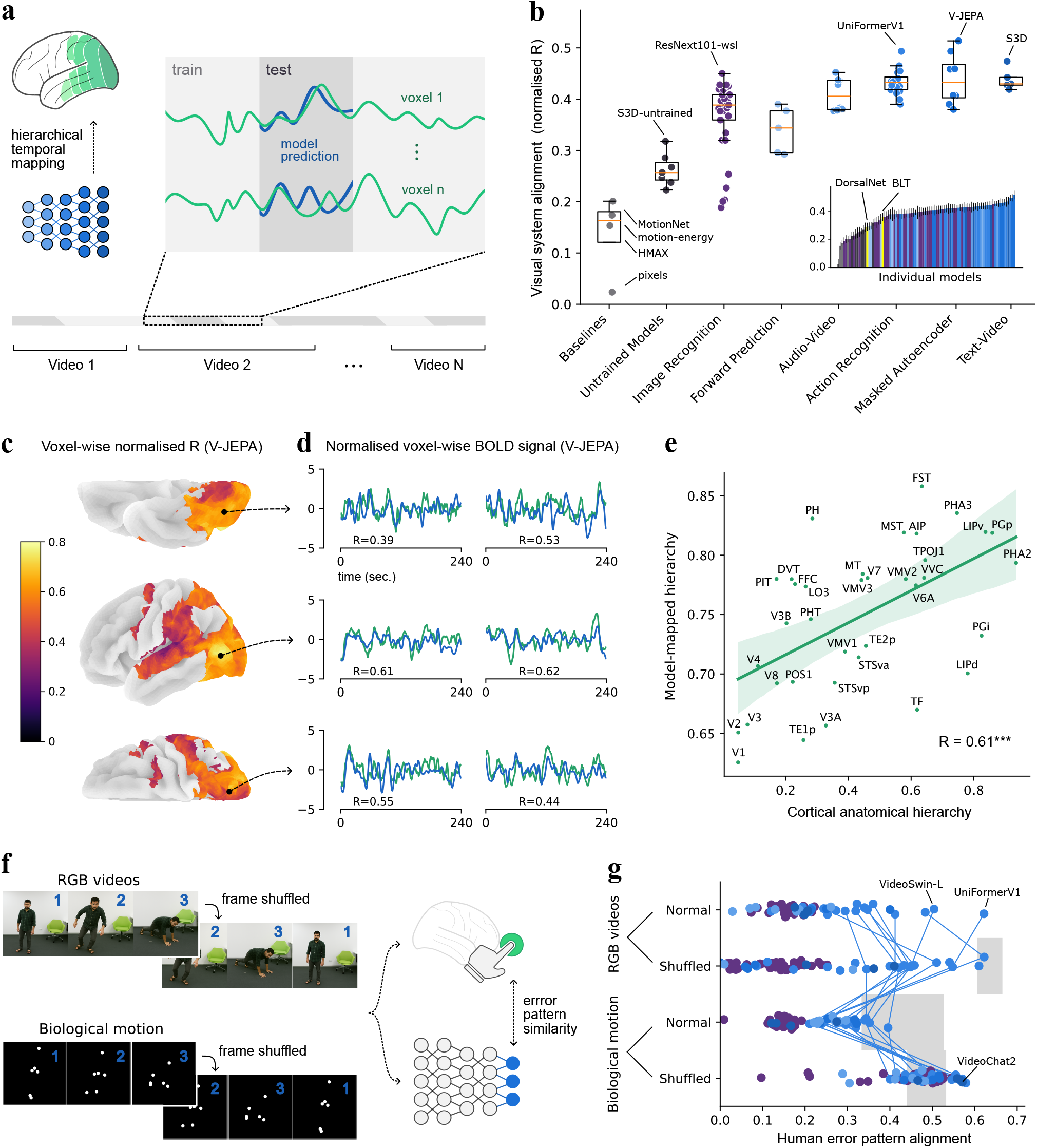
Predictive models of dynamic vision. **a. Neural alignment:** we map layer-wise representations from stimulus-computable models to dynamic neural signals (fMRI voxel time series) and evaluate similarity across time. **b. Dynamic models are most brain-like:** many dynamic task-optimised models (blue shades) outperform baselines and static models (purple). Inset: individual model scores (details in SI. A.3). **c. Alignment scores of the best model:** normalised fMRI predictivity of V-JEPA on a 3D brain atlas from ventral, symmetrically flipped lateral, and dorsal views. Showing the right hemisphere, similar patterns are observed in the left hemisphere (Fig. S4). **d. Second-by-second predictions:** predicted fMRI responses from V-JEPA for held-out 240-second video clips (blue) compared to ground truth (green) from the three views in panel **c. e. Model hierarchy matches cortical hierarchy:** layer assignments averaged across the 20 best-aligned dynamic models strongly correlate with the anatomical hierarchy derived from *Rolls et al*. [142] (*p<* 0.001; regions from *Glasser et al*. [56]; Methods B.3 and SI A.6). **f. Behavioural alignment:** we compare action recognition judgements on RGB or biological motion videos (with either shuffled or intact frame order) between humans and models, based on recognition error patterns. **g. Models predict human video error patterns:** model alignment to human errors (dots). Grey shaded areas show inter-subject consistency (95% CI). Lines connect individual top models across conditions.

To investigate which computational objectives shape brain-like representations, we further evaluate the performance of ANNs on 10 ecological tasks (Fig. 1a right). These include object recognition—long a focus of prior work [174, 147, 99]—as well as dynamic tasks [104, 175, 64, 150] like action recognition [143, 154, 7, 129] and social understanding [52]. We evaluate voxel-level task processing and provide evidence that a single functional objective underlies the formation of representations in the visual system.

### Task-optimised dynamic neural networks predict cortical dynamics

We first test if models can predict the brain activity elicited by video viewing. Candidate ANNs are sourced from six pretraining classes: image recognition, forward prediction, audio-video correspondence, action recognition, masked autoencoding, and text-video models. All task-optimised model families accurately predict cortical dynamics, up to 0.51 correlation with visually selective voxels across the brain (Fig. 2b, coloured dots).

#### Task-optimised models outperform baseline and untrained models

Compared to baselines—such as a classic neuroscience model [139, HMAX], spatiotemporal filtering [124, motion-energy], and simple dynamic models [116, 138, MotionNet]), models optimised for image or video tasks achieve significantly higher brain alignment (mean normalised correlation 0.40 vs. best-performing baseline 0.20; *p<* 0.001). Trained models outperform their untrained counterparts (*p* = 0.011; SI A.2), which themselves outperform all baseline models (*p<* 0.001, Fig. 2b). This suggests that a combination of structural priors and task optimisation is critical for brain-like representations. Notably, these models are optimised for a single objective and yet learn representations that predict activity across the visual cortex.

#### Dynamic models outperform static models

Many dynamic model families show strong brain alignment, significantly outperforming static object recognition models (action recognition: *p<* 0.001; masked autoencoders: *p* = 0.015; text-video models: *p* = 0.002; Fig. 2b).

The most brain-aligned model identified here, V-JEPA [5], learns representations by reconstructing missing features in video segments – achieving a normalised correlation to brain data of 0.51, with reliable predictions across ventral, lateral, and dorsal areas (Fig. 2c,d). Predictivity decreases in regions adjacent to the auditory cortex (SI A.1&A.5), which can be attributed to the unimodal nature of the model.

#### Model-derived hierarchy matches empirical connectivity

During alignment, we map different layers of the neural network to distinct brain regions based solely on functional correspondence, without using anatomical constraints. Remarkably, the cortical hierarchy emerges from this functional mapping. We quantify this hierarchy by averaging the relative layer positions assigned to each brain region across the 20 best-aligned dynamic models, yielding high split-half consistency (*r* = 0.84). Indeed, this model hierarchy aligns closely (*R* = 0.61, *p<* 0.001, Fig. 2e) with cortical connectivity in the brain [142]. For example, the layer best aligned with V1 appears early in the model, while higher-level regions such as ventral VVC and dorsal MST correspond to later layers (SI A.6).

#### Dynamic models make human-like decisions in action recognition

Beyond a characterisation of the neural substrate underlying dynamic vision, we link our findings to visual behaviours. Using a large-scale dataset of indoor actions [66], we compare action recognition decisions from models and humans across two presentation formats (RGB and biological motion) and two temporal ordering (normal and shuffled) conditions (Fig. 2f; Methods B.4).

To test behavioural alignment, we examine whether the models exhibit similar error patterns to those of humans (with accuracy results reported in SI A.7). Specifically, we assess the similarity of object-level confusion between humans and models, defined as the probability of misrecognising one action as another [134]. Models closely align with human error patterns, with many models approaching the inter-subject consistency observed among human participants (consistency is undefined for the first condition due to insufficient errors in RGB video trials; Fig. 2g; Methods B.4).

### Two principal computations explain model-brain alignment

The strong alignment of task-optimised neural networks with brain activity prompts the question: *what underlies this alignment?* Evidence from recent studies suggests that brain alignment in individual streams is closely tied to task performance, particularly on ecologically relevant behaviours [174, 147, 81, 146, 122, 109].

Unlike prior work, our study analyses the entire visual system and thus calls for evaluating a broad range of candidate cognitive tasks and their associations with brain regions. Our framework is scalable: via stimulus-computable models, we can directly test arbitrarily many task and stimulus conditions, focusing our investigation on the most striking effects. We examine ten candidate cognitive tasks spanning static, dynamic, and hybrid information processing—including object recognition, action recognition, and social understanding (see Figs. 3a and S10 for sample stimuli). To also test the capacity of processing appearance-free motion (*i*.*e*., dynamics without colour or texture cues), we include three tasks for direction discrimination, motion-only action recognition, and biological motion recognition. To assess a model’s capacity to support a task independently of its pretraining objective, we evaluate performance using a linear decoding probe (Methods B.5).

**Figure 3.**
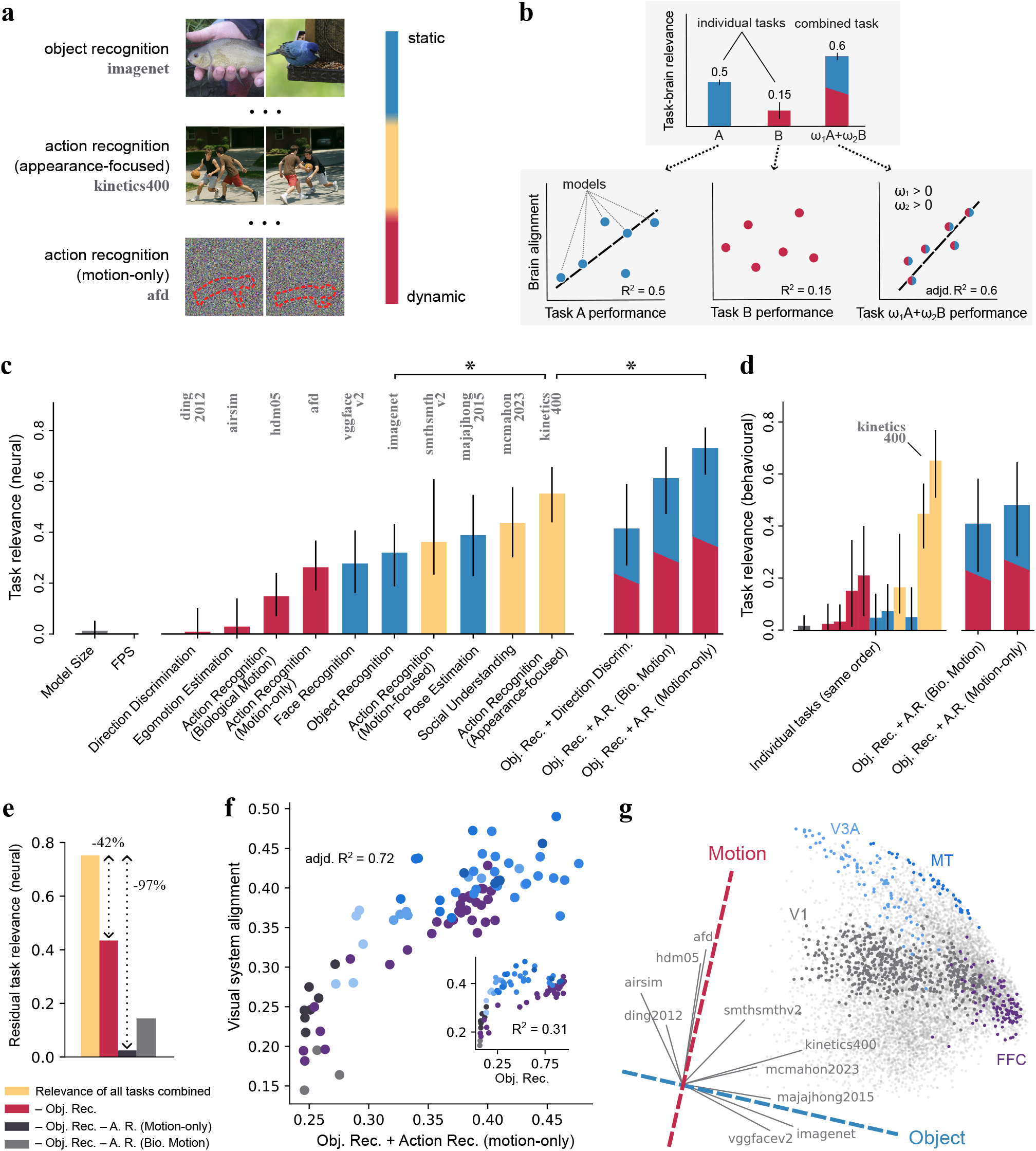
Object and appearance-free motion recognition jointly explain representational alignment. **a. Diversity of tasks:** We evaluate models’ cognitive capacities on a broad range of tasks (complete task suite in SI B.1). Static tasks test for image-based object recognition, while purely dynamic tasks are devoid of appearance cues (motion highlighted with dashed red outlines). **b. Testing cognitive tasks for their relevance to brain function:** To test the influence of multiple tasks on shaping brain-like representations, we assess individual and joint *task relevance* as the correlation between models’ task performance and their brain alignment (assessed via *R*^2^; adjusted *R*^2^ for multivariate regressions). **c. Neural task relevance:** Task relevance for brain alignment of controls (grey), individual tasks (single colours), and joint tasks (mixed colours). Action recognition is individually most predictive; combined object and motion recognition explain most variance overall (SI B.2). **d. Behavioural task relevance:** same as in **c**, but evaluated with respect to behavioural alignment. **e. Residual relevance:** relevance of all tasks combined after regressing out the contribution of certain tasks. Joint object and motion recognition explain away most of the relevance (voxel-wise results in Fig. S13). **f. Object and motion task performance predicts brain alignment:** correlation between object + motion-only action recognition and brain alignment. The correlation of object recognition alone (inset) is significantly lower (*p<* 0.01). Model groups coloured as in Fig. 2b. Regional results in SI B.4 **g. Task geometry in visually driven cortex:** the 2D eigenspace captures 91% of the variance in task relevance across visually driven voxels in the brain. This space is spanned by two orthogonal basis directions: object and motion recognition.

#### Naturalistic hybrid tasks individually yield highest relevance

We identify tasks where models’ performance is most correlated with their brain alignment (“*task relevance*”; Fig. 3b). A high correlation suggests that optimising for the task leads to brain-like representations. Appearance-focused action recognition (`kinetics400`) emerges as the single most *relevant* task predicting models’ overall brain alignment (*R*^2^ = 55%; Fig. 3c). It significantly surpasses object recognition (`imagenet2012`; *R*^2^ = 32%) which has been closely tied to the primate visual ventral stream [174, 147]. Other tasks that involve processing naturalistic dynamics also show high *task relevance*, including social understanding and motion-focused action recognition. Controls such as model size and frames-per-second (FPS) are not predictive of brain alignment.

#### Two tasks explain the diversity of cognitive processing

Following the intuition that the brain’s neural code might be driven by multiple cognitive demands, we sought to explain model-to-brain alignment as a combination of task performances. We introduce a novel *joint task relevance*, which weighs each task to maximise correlation with brain alignment (Fig. 3b; Methods B.6). Applied to all candidate tasks, object and appearance-free motion recognition emerge as the combination that explains the most variance in brain alignment (*adjd. R*^2^ = 73%, Fig. 3c right), significantly outperforming any individual task (*p <* 0.05, FDR-corrected; Fig. 3f). Importantly, this combination explains away all other tasks (Fig. 3e), while object recognition alone only accounts for 42% of the contribution (SI B.2). The same combination also shows strong relevance to behavioural alignment (*adjd. R*^2^ = 46%, Fig. 3d; SI B.7).

#### Two tasks explain the complexity of neural and behavioural representations

Since object and appearance-free motion recognition together explain nearly all the variance in neural alignment, we asked: *do these two components span the representational organisation across the visual cortex?* We computed voxel-level task relevance across the 10 cognitive tasks, yielding a 10-dimensional relevance profile for each voxel. Fig. 3g presents the two-dimensional eigenspace projection of these profiles, capturing 91% of the total variance (44% and 47% on the two principal dimensions). The projection can be spanned by two orthogonal axes for motion and object processing, with motion- and object-focused tasks near their respective directions, and naturalistic dynamic tasks centrally, reflecting hybrid processing (grey arrows in Fig. 3g). Patterns also emerge for brain regions: dorsal regions such as V3A and MT show strong motion bias, ventral regions like FFC favour object processing, and early visual areas (*e*.*g*., V1) show a balanced profile, integrating both components.

We examine regions across the ventral and dorsal visual streams, including the dorso-dorsal and ventro-dorsal subdivisions. The combined task explains the majority of the variance in brain alignment across nearly all regions (Fig. 4a), with distinct biases toward either object or motion processing in each region.

**Figure 4.**
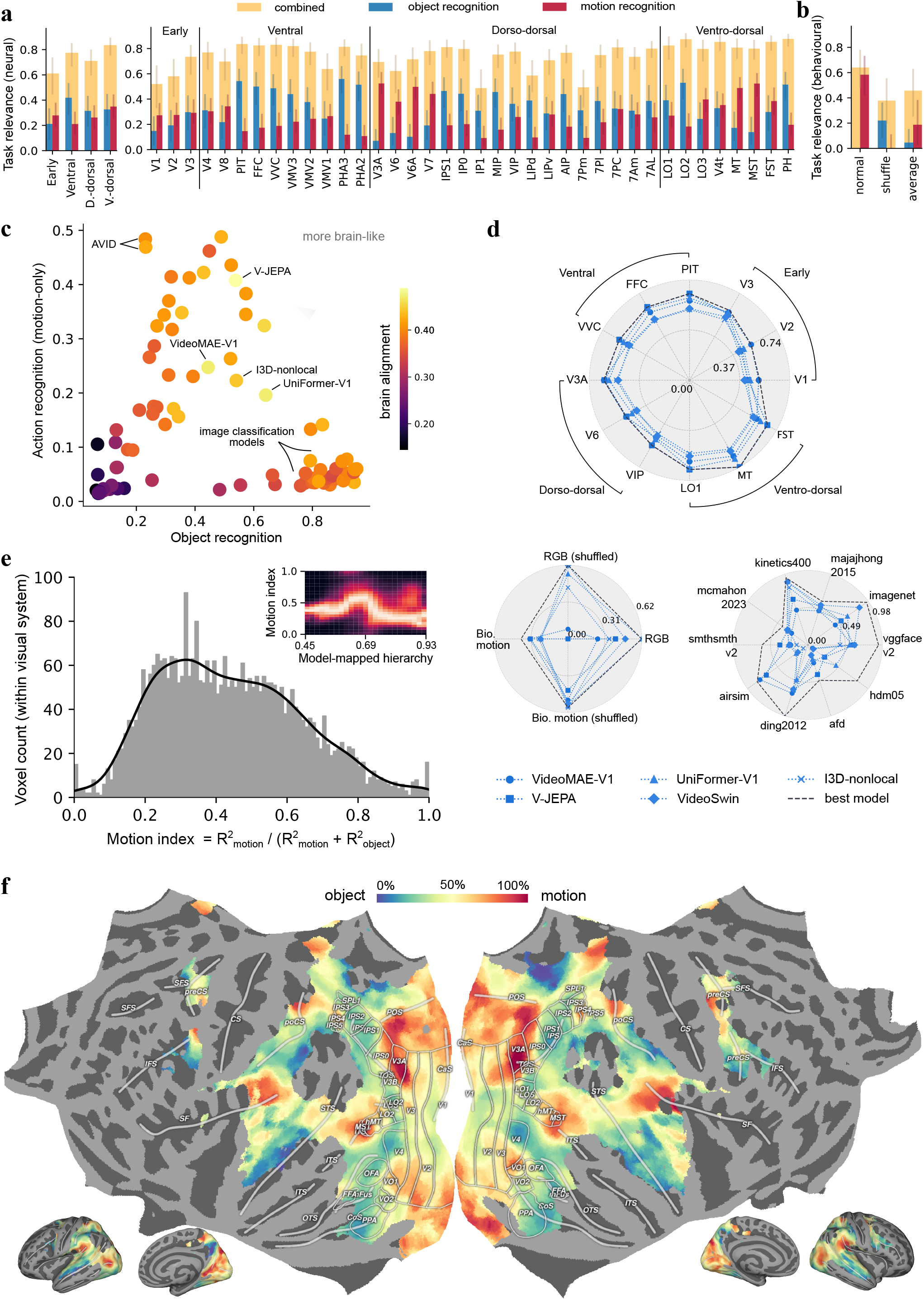
Diverse visual brain representations originate from a single objective. **a. Joint object and motion recognition are most relevant for all visual ROIs:** task relevance of object recognition, motion recognition, and their combination to regions of interest (ROIs; SI B.3) along ventral and dorsal pathways. **b. Joint object and motion recognition are most relevant for behaviours across video conditions:** task relevance of action recognition behaviours in normal videos, temporally shuffled videos, and their average. **c. Single objectives jointly capture object and motion recognition for maximum brain alignment**: task performance on object recognition versus motion-only action recognition. Multiple single-objective models successfully implement both capacities, resulting in higher brain alignment. **d. Single objectives yield consistently high neural alignment, behavioural alignment, and task performance:** evaluation of models trained on single objectives across alignment and performance measures. Each connected line is one model. **e. Smooth distribution of motion index across the visual system:** motion index reflects the proportion of motion versus object processing within each voxel. The inset shows the distribution across the full visual hierarchy (filtered from Fig. S26). **f. Cortical map of motion index:** brain-wide motion index map (for voxels with at least 40% inter-subject consistency; Methods B.1). Hybrid processing and smooth transitions are observed in all visual streams.

In terms of behavioural relevance, motion recognition primarily supports decisions with natural videos, whereas object processing dominates under temporally shuffled videos, reflecting two distinct processing mechanisms under these conditions (Fig. 4b).

### Brain-like representations emerge from single objectives

Representations across visual cortex can be explained by two tasks: object recognition and appearance-free motion recognition. We next examine how these two computations are distributed across the brain and present evidence that brain-like representations are ultimately implemented by a single unified objective.

#### Distributed computations across visual brain regions

For all visual brain regions, hybrid processing dominates, with a smooth transition of preferential object and motion processing along visual streams (Fig. 4a right). For example, mid-level regions in both dorsal subdivisions, such as V6A and MT, are biased toward motion processing, but both converge to mixed processing at later stages. Processing of object and motion information is nearly evenly mixed across visual pathways although the ventral stream shows a notable bias toward object recognition (Fig. 4a left). While the classic view of distinct pathways with distinct functions would have suggested strongly biased functional preferences such as object recognition in the ventral pathway and motion recognition in the dorsal pathway, the ubiquitous dominance of hybrid processing and smooth gradient across regions suggest representations might instead be shaped by a single unified objective.

#### Single-objective models achieve brain-like representational diversity

Models pre-trained on single objectives demonstrate high alignment with the entire visual system (Fig. 2b). The best models achieve high brain alignment not by improving only object or motion recognition, but via an objective that implicitly optimises both (Fig. 4c). Indeed, across visual regions, behaviours, and cognitive tasks, several single-objective models exhibit consistently high alignment and performance (Fig. 4d). Brain-like representations thus can in fact emerge from a unified objective such as action recognition or masked autoencoding, which all implement a form of world modelling.

#### Absence of multimodality in voxel task relevances suggests a unified functional objective

Beyond regional divisions (Fig. 4a), we investigate the detailed differences in motion and object preferences on a voxel-by-voxel basis. We define a *motion index* to quantify the proportion of motion recognition relevance relative to object recognition relevance. The distribution of motion index values is smooth across voxels (Fig. 4e): rather than exhibiting a bimodal structure indicative of distinct functional clusters, the distribution is approximately unimodal (*p* = 0.993, Hartigan’s dip test [105]). This pattern is consistent across the visual hierarchy (inset in Fig.4e; Fig. S26).

Throughout cortex, motion and object preferences across voxels vary smoothly (Fig. 4f), suggesting that these functions are not strictly segregated but rather closely intertwined. Functional organisation in the cortex following a topographic map is a well-established phenomenon [72, 74, 67, 169, 108] and helps explain the bifurcation of specialisation despite a single overarching objective. The distribution of voxel-by-voxel task relevances exhibits spatial clustering which follows a topographic organization – with smooth transitions throughout rather than a strict division into functional streams.

### Task-based functional localisation of dynamic vision networks

The motion index map (Fig. 4f) reveals cortical structures corresponding to prior neuroscience findings on object and motion processing in both visual streams. Here, we interpret these structures and link them to behaviour, offering strong validation for our methodology.

#### Unification of fragmented experimental findings

The division between object and motion processing in the motion index map reveals a functional organisation that aligns with prior neuroscience findings, offering a unifying perspective on fragmented observations across visual streams. Fig. 5a shows that motion processing aligns with visual cortex eccentricity: peripheral regions such as MT and peripheral EVC are more involved in motion processing than foveal EVC, consistent with known retinotopic organisation [15, 170, 69]. Voxels with a bias toward object processing (motion index *<* 45%) match regions identified in the shape processing network (Fig. 5b), including LO, PHAs, fusiform gyrus, and IPS [47, 39]. This finding is consistent with recent observations that image classification models explain activity not only in the ventral stream but also in dorsal areas [45] (Contrasting the task relevance of face and action recognition, our computational analysis also rediscovers the face processing network, Fig. S18). Voxels biased toward motion processing (motion index *>* 55%) on the other hand correspond to regions responsive to optic flow and implicated in egomotion encoding (Fig. 5c) [20]. These regions include V1–V3 in early visual cortex; MT and MST in the ventro-dorsal stream; and dorso-dorsal areas such as V3A, V6, and VIP. They also include the parieto-insular cortex (PIC), located between the inferior parietal lobule and the auditory cortex [46]. The motion features encoded across these regions increase in complexity along the visual hierarchy (SI B.5). The spatial functional gradient suggests potential pathways linking these motion-sensitive regions, which we explore next.

**Figure 5.**
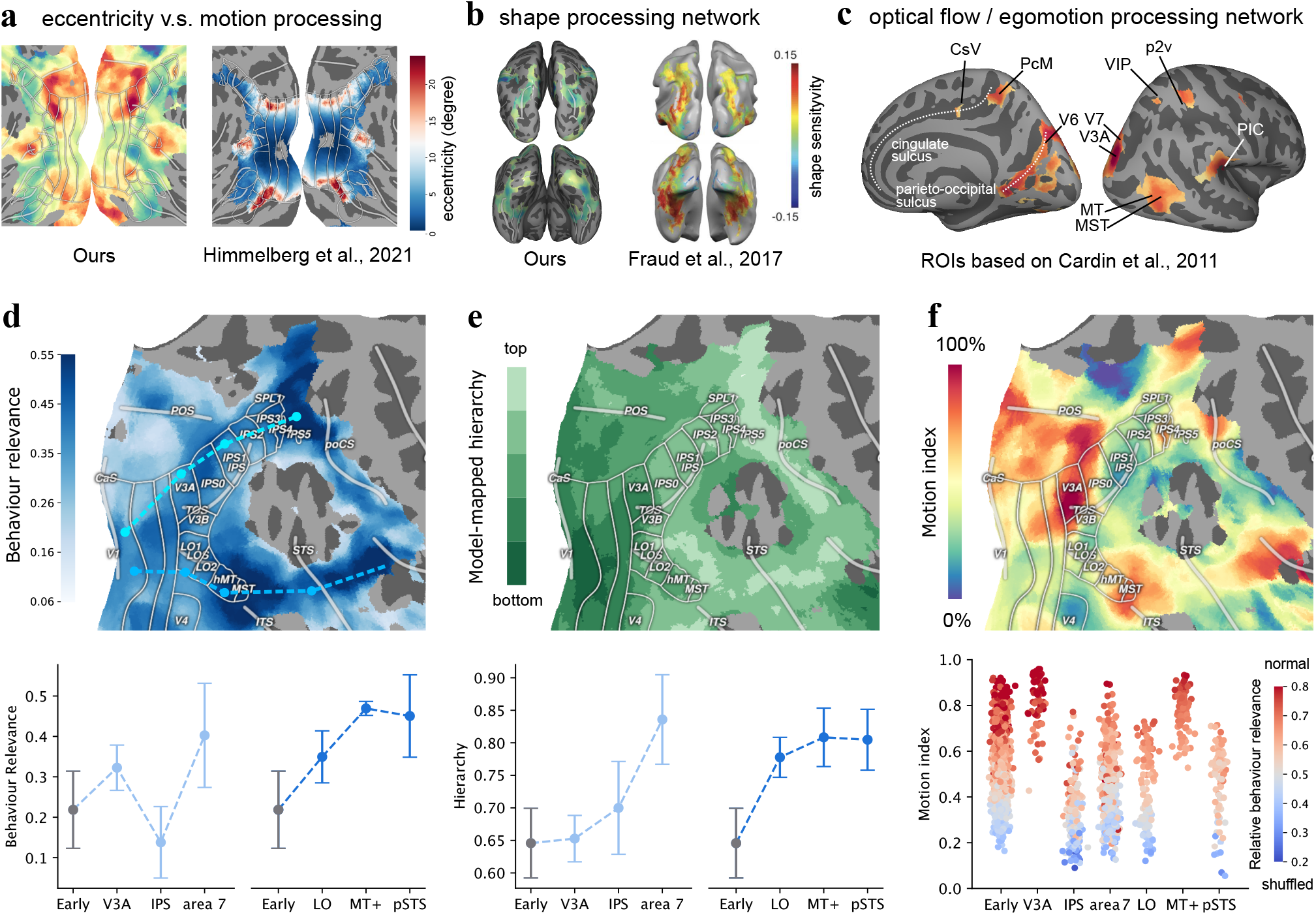
**(a-c). Recapitulation of known brain structures:** *a*. motion index in early visual cortex (EVC) compared to EVC eccentricity [69]; *b*. object-biased areas of our map (*<* 45% motion index) compared to the shape processing network [47]; motion-biased areas of our map (*>* 55% motion index) cover the optical flow/egomotion processing network [20]. **(d-f). Computational characterisation of action understanding pathways:** *d*. behavioural relevance increases along dorso-dorsal and ventro-dorsal streams (full map in Fig. S17). *e*. model-mapped hierarchy increases along dorso-dorsal and ventro-dorsal streams (full map in Fig. S6). *f*. hybrid processing occurs across many regions along the streams; a higher motion index corresponds to greater behavioural relevance under normal temporal video order, and a higher object index for shuffled conditions.

#### Emerging pathways of action understanding

The motion index map suggests processing that starts from initial broad functional processing in early visual cortex and then, through two subdivisions of the dorsal stream, converges to a high-level hybrid nature of representations. We here link this functional analysis to behavioural relevance and computational hierarchy (Fig. 5d,e) for a detailed characterisation of action understanding pathways. Tracing regions along the dorso-dorsal (V3A, IPS, area 7) and ventro-dorsal (LO, MT+, pSTS) streams (blue dash lines in Fig. 5d) reveals increasing behavioural relevance (Fig. 5d, bottom) and computational hierarchy (Fig. 5e). With hybrid object and motion processing across regions, intra-regional subdivisions seem specialised for different conditions of temporal integration: voxels with a higher motion processing index are more behaviourally relevant for normal videos while voxels with an object bias relate more closely to behavioural choices for shuffled videos (Fig. 5f). On the ventro-dorsal stream, the relevant regions seem to extend beyond pSTS and go into the posterior Sylvian fissure, which is adjacent to the posterior insula cortex (PIC). While previous studies have examined this region in the context of egomotion and vestibular processing [46, 132, 20], its potential role within an action understanding pathway as suggested here has been largely overlooked [140, 142, 11, 171, 26].

## Discussion

### Summary of key results

Our results show that specific artificial neural network models can predict neural and behavioural responses to dynamic visual input with high accuracy: The best models are dynamic in nature, organised in a way that resembles the cortical hierarchy, and optimised to understand the visual world. Dynamic visual representations are computationally organised in a space spanned by two principal axes, which are respectively in support of object and appearance-free motion recognition. While these two tasks preferentially map onto classic ventral and dorsal streams, computations throughout visual brain regions are fundamentally distributed and hybrid. Our task-based functional localisation shows that the organisation of representations on tissue follows a smooth interpolation between object and motion processing, exhibiting an overall unimodal distribution. Furthermore, models optimised for a single objective yield representations with high alignment to all brain regions, reproduce key experimental results, and exhibit consistently high behavioural accuracy. Together, these findings point to a unified functional objective shaping the visual system.

### Perceptual representations are driven by a single objective

Building on decades of research in visual neuroscience and connecting to advances in AI allows us to address the fundamental question of *how* representations are shaped in the brain. Partial accounts of visual cortex representations had pointed to object recognition for the visual ventral stream [174, 147, 17, 89, 84, 123, 78, 148, 25], with recently increasing studies on non-ventral areas [116, 23, 3, 44, 143, 157, 24]. We here aim to provide a holistic view of human dynamic perception by identifying models and computational principles for the entire visual cortex. Two core findings emerged, both supporting the idea that the human visual system is optimised for a single ‘world understanding’ objective.

First, optimising for a single objective yields the most brain-like models. The exact details of the optimisation procedure vary – from predicting masked parts in a video to recognising actions and events – but all attempt to understand the dynamics in the visual world. The brain alignment of models trained with a single world modelling objective seems to continue to scale with better and better models, which might overcome the saturation of models trained with more narrow objectives [57, 99, 147]. After training, the emergent representations in these models are highly diverse: they align with areas from early visual cortex to high-level regions, spanning both ventral (*e*.*g*., FFC) and dorsal (*e*.*g*., MT, VIP) streams. Meanwhile, they also support various action recognition behaviours and high performance across cognitive tasks.

Second, representations in visual cortex are organised in a smooth and hybrid manner. When we analysed the task relevance of visual regions, we found that a hybrid combination of object and appearance-free motion recognition is most explanatory and explains away all other tasks; a similar pattern holds for visual behaviours. Meanwhile, voxel-wise organisation of these computations is smooth: the vast majority of voxels support both object and motion recognition, and the overall distribution is unimodal across the visual hierarchy. If representations were shaped by multiple objectives, we would expect a multimodal distribution that is centred around two or more tasks with many visual regions being entirely dominated by one task; but we only found evidence to the contrary.

An intriguing possibility is thus that both the biological visual system and successful vision models are ultimately optimised to understand the visual world. The need to be competent across a wide spectrum of perceptual tasks may drive both systems toward convergent solutions, a trend likewise increasingly emphasised in recent artificial intelligence literature [73, 13, 97]. To engage on this question, our multi-task relevance analysis is applicable to arbitrary species and brain regions, enabling tests of how diverse tasks drive neural processing and might converge on a shared set of core features.

### Organisation of representations into pathways

Beyond the developmental origin of visual representations, we also investigated the resulting cortical organisation. Ever since the proposal of the two-visual-stream framework, a large body of work has sought to refine and extend it [119, 61, 114, 140, 171, 110, 85, 142, 11]. Stream-based theories of visual processing have become predominant in the literature and remain canonical in shaping current understanding of the visual cortex, with only limited challenges to this view [29, 145]. However, these descriptive models are established primarily by the qualitative aggregation of fragmented observations from functional neuroimaging, lesion, intervention, and neuroanatomical studies [171, 140, 11], with individual pieces of evidence often limited to specific regions and experimental conditions. In contrast, our model-based framework integrates a wide range of neural and behavioural datasets and, critically, enables quantitative predictions for any novel stimuli. At the same time, such unifying models enable analyses with system-level breadth alongside voxel-level precision. These properties enable us to investigate visual processing at the scale of the entire cortex, spanning the regime of multiple visual streams.

Here, we connect the representations of brain-like neural network models to different cognitive tasks by asking: *Which* tasks drive the alignment to *which* parcels in the brain? We developed a novel method (Fig. 1a & Fig. 3b) which jointly considers the relevance of multiple tasks to cortical regions, moving beyond previous analyses on individual tasks [174, 81, 146, 122, 109]. Our analysis offers a potential nuanced resolution on visual stream theories: in line with classic ventral-dorsal literature, representations *preferentially* support object over motion recognition and vice versa — but neural mechanisms and their resulting representations appear consistently hybrid due to their shared single-objective origin. Our task-based functional localisation offers a new method for mapping the entire processing hierarchy (SI B.8) and further characterises these hybrid representations at voxel-level resolution. The localisation map reproduces a number of previously separate experimental findings on distributed computations across visual streams: eccentricity [69] and motion processing correspond such that more peripheral regions of early visual cortex are more involved in motion than foveal parts; the shape processing network [47] corresponds with object-biased voxels; and regions implicated in optic flow and egomotion [20] are closely linked to voxels biased toward motion processing. These observations validate our mapping technique. Furthermore, by integrating them into a unified map, we reveal the continuum of object versus motion processing across the entire hierarchy of the visual system, beyond specific regions and experimental contexts.

A potential explanation for this hybrid and continuous organisation is that the unified functional objective of the brain drives feature encoding, while cortical wiring constrains how they are spatially arranged, resulting in a topographically organised functional architecture [108, 34, 12, 82, 136, 30]. This would reconcile with the evidence against strict separation between visual streams [145, 141], and integrate emerging proposals of new pathways [171, 60, 85].

Our map further suggests that the ventro-dorsal stream supporting action understanding extends into regions near the posterior insular cortex, which are associated with multimodal vestibular-visual and auditory-visual (human planum temporale) processing [27, 65, 137]. This observation aligns with anatomical connectivity studies [65, 137] and highlights the multimodal nature of action understanding and, more broadly, world understanding.

### Limitations and future directions

While this and related studies aim to understand the brain in computational terms, adequate brain recordings remain essential for robust findings. High-quality recordings are hard to come by, especially at a scale sufficient for model testing, and most datasets in the field are not collected with model comparisons in mind in the first place. To integrate the 35.7 hours of recordings from 5 different fMRI datasets, we chose to map all the recordings of different subjects onto the *MNI* standard space and focus on group-level analysis only. This likely leads to lower internal consistency for signals in higher-level areas that perform long-term integration and memory [144], such as the prefrontal cortex, medial temporal gyrus, and angular gyrus. The fMRI BOLD signal is also relatively coarse, especially in the time dimension, and prevents us from a finer-grained investigation on millisecond-resolution dynamics. High-resolution datasets at scale with high repetition counts would be ideal, but likely require larger-scale experimental initiatives. To maximally discern between alternative model candidates, we believe including models in experimental designs is a vital step [58, 71].

The discoveries from this work enable exciting future directions. Due to the solid alignment of the best models, they can now be used to guide experiments. For instance, past studies have shown that model-selected stimuli can control neural activity [8, 133, 166, 163]. The models identified here potentially enable such model-guided control for temporal stimuli. Further, the best models can be analysed to make sense of their inner workings. Interpreting the function of computational models is a lot more accessible and efficient than biological experiments due to the complete availability of structure, functional connections, and the ability to perform targeted ablations.

The best models identified in this work do not hit the noise ceiling of the data. How can we further improve models of dynamic vision? With the hierarchical motion network culminating in areas associated with multimodal processing, including other sensory modalities in computational models might be a required next step. It is conceivable that a world modelling objective underlies not just visual cortex but sensory cortices more broadly and the division between “domain-specific” areas might be much smoother, as we found here for areas in the visual system. In this case, building models that comprehensively understand the world across modalities might not only lead to a more unified model of sensory processing in the brain, but also improve the alignment to individual sensory modalities.

### Conclusions

Taken together, our findings suggest that temporal AI vision models serve as viable hypotheses for how the human brain processes visual input in the service of understanding the world. This computational framework together with a novel multi-task relevance method explains how the visual system might be optimised for a single objective — and yet organises its representations along principal axes of object and motion recognition. We thus offer a potential resolution to previously conflicting views on visual streams where the classic dual-stream organisation emerges as a slight bias of a unifying goal, but with an overall smooth and inherently hybrid organisation of visual cortex in the service of integrating dynamic information.

## Methods

### A. Experimental model and study participant details

We use fMRI recordings from five different studies, three [141, 9, 80] of which use movies lasting 6 to 10 minutes as stimuli, and two [111, 90] use short video clips of duration 3 seconds as stimuli. We also use macaque electrophysiological recordings from four visual areas, collected across two separate datasets with distinct image stimuli comprising objects, textures, and noise patterns [177, 107] (SI A.4). Additionally, we analyse a behavioural dataset involving video viewing and action recognition [66], in which human psychophysics experiments were conducted on Amazon Mechanical Turk. The experimental background can be found in the respective publications.

### B. Method details

#### B.1. Neural and behavioural datasets

##### Neural datasets of movie/video watching

We analyse a battery of movie- or video-watching datasets for studying the temporal neural dynamics in the neocortex with pseudo-natural visual stimuli. These datasets include a rich diversity of dynamic visual information and a high number of subject hours. *Berezutskaya et al. 2021* [9] contains fMRI recordings from 30 participants, all of whom performed a passive movie-watching task on a short 6.5-minute audiovisual film. *Keles et al. 2024* [80] includes fMRI and eye-tracking data from 24 participants during an 8-minute excerpt from a movie, after which they were asked to perform a recognition memory test. *Sava-Segal et al. 2023* [144] includes fMRI data from 43 subjects who watched four movies, all of which lasted for about 8 minutes and were followed by a set of tasks to probe the participants’ interpretations and reactions. Although we focus only on modelling vision, the above three datasets all used audio-visual movie stimuli. In contrast, the following two use purely visual stimuli: in *Lahner et al. 2024* [90], 10 subjects viewed 1,000 3-second videos with 1-second intertrial intervals and with fixation. They were also presented with null trials 25% of the time during which they were instructed to press a button. Finally, *McMahon et al. (2024)* [111] have four participants who freely viewed 250 3-second videos featuring two actors (with or without interaction between the two actors). The statistics of these datasets are summarised in SI A.1.

##### MNI space registration and fsaverage extraction

In this study, all fMRI volumes are registered to the *MNI152NLin2009cAsym* template [36] (hereafter abbreviated as *MNI*) either by the original research work or by us. We refer the readers to the individual papers [9, 80, 90, 111, 144] for their detailed preprocessing steps. Then, the cortical signals in the *fsaverage5* mesh space are extracted for the subsequent analyses. For registered volumes, we use *nilearn* [1] to extract signals from the *fsaverage5* mesh space. For non-registered volumes, we directly run *fMRIprep* [36] with default settings and specify the output space to be *fsaverage5*. In the latter process, we also regress out the principal components of estimated nuance factors, retaining the top 95% variance. For all volumes in each dataset used in the study, we average across subjects and repetitions, focusing solely on the group-level analysis (SI A.1). For visualisation purposes (with pycortex [50]; Fig. 4f and Fig. 5), we upsample the data on the *fsaverage5* mesh to the *fsaverage* mesh using a 10-nearest neighbours (KNN) averaging procedure. The locations of the vertices from both meshes in the 3D *MNI* space were taken from *nilearn*.

##### Normalisation over time

To utilise data from all sources simultaneously, we simply standardise the fMRI signals to have a mean of zero and a variance of one across time, separately for each dataset. This normalisation disregards many differences between the datasets, such as the MRI device, original spatial resolution, thermal noise, *etc*., making the cross-dataset predictions of fMRI signals particularly challenging. However, this coarse normalisation is sufficient to produce reasonable predictive performance from the models studied in the paper. We plan to investigate finer-grained normalisation in future work.

##### Internal consistency computation

The neural signals are inherently noisy. To fairly assess the predictions for these signals, we first estimate the internal consistency for each dataset at the group level. Specifically, denote *x*_*v,t,s,r*_ as the fMRI BOLD signal at voxel *v* and time *t* for subject *s* at the *r*-th repetition, the internal consistency *c*_*v*_ for this voxel was computed as:

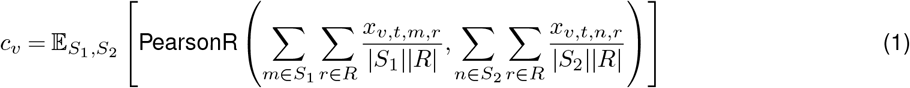

where *S*_1_ and *S*_2_ are even splits of the whole set of subjects, *R* is the total number of repetitions, and *PearsonR* is the Pearson correlation over time *t*. The expectation is estimated using 10 random samples of the splits. This internal consistency is then corrected using the Spearman-Brown correction for two splits: 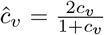. This voxel-wise measure indicates the ceiling of Pearson correlation a predicted signal can have with the averaged group-level signal. For all voxel-wise analyses, we use this measure to normalise model predictivity. We estimate the normalised ROI-wise (and whole-brain) predictivity by averaging the normalised voxel-wise predictivity within each ROI (or the whole brain). All of these analyses are restricted to the voxels with an internal consistency greater than 0.4 (Fig. S27). We choose this threshold due to the fact that it covers most of the action observation network [54], which has been implicated in processing dynamic event information while ensuring overall signal reliability.

##### Behavioural datasets

We use a subset of a recent large-scale behavioural dataset examining participants’ action recognition with RGB and biological motion videos [66]. The biological motion presentation focuses exclusively on the key joints of the actors in the videos. This minimalist representation, when contrasted with the detailed RGB videos, provides valuable insights into the motion aspects of human action recognition. Furthermore, the videos are shown in two distinct temporal ordering conditions: normal, where the frames are displayed in their natural chronological sequence; and shuffled, in which the frames are randomly shuffled in time. These conditions allow us to examine the temporal integration over time. The human psychophysics experiments were performed on Amazon Mechanical Turk with 90 recruited participants. Participants were presented with a video (a few seconds long) and a list of ten options, then asked to complete a forced-choice 1-out-of-10 test to identify the action in the video. For the subset we analyse, each of the four conditions includes 600 trials, resulting in a total of 2,400 trials across all conditions. We refer readers to the original paper [66] for details on experimental design, data filtering, and stimuli selection. Beyond the stimuli used in the behavioural experiment, since this dataset itself is a subset of the NTU RGB+D 120 dataset [100], we are able to collect additional data with ground-truth labels across different conditions. These data are used for training the behavioural decoder trained on model outputs, as will be in Methods B.4. Sample images shown in Fig. 2 are drawn from the same dataset. We regenerated them using ChatGPT [127] to ensure anonymity.

### B.2. Activation extraction from deep neural networks

#### Model groups

We evaluate 92 pretrained deep learning models across many architectures (convolutional neural networks, attention-based transformer models, recurrent models, and their hybrids) that are chosen to represent a diverse range of training objective designs. They include: 1. baselines models based on traditional computational modelling, including “pixels”, HMAX [139], motion-energy [124], and MotionNet [138]; 2. the forward-prediction models that were trained to predict the next frame in a video sequence, including PredRNN [135], ConvLSTM [151], SimVP [51], and MIM [168] as implemented by [155]; 3. the audio-video models trained to align audio representation with corresponding video representations, including SeLaVi [2] and AVID [120]; 4. the action recognition models trained to recognise human actions, including convolution-based [161, 172, 21, 41, 40, 10] and transformer-based models [38, 96, 102, 93, 92]; 5. the mask autoencoding models trained to reconstruct missing spatiotemporal patches in videos in the pixel or latent representation space, based on transformer models [159, 167, 5, 42]; 6. the text-video models trained on text-video contrastive matching (S3D-Howto100M [113, 112]) or various video-conditioned text generation tasks such as VideoChat [91] and Video-LLaVA [98]; 7. the image recognition models trained to classify objects, including those identified as among the most brain-like models (ResNext [106], CORnet [89], and VOneNet [28]), AlexNet trained on ImageNet or Stylized ImageNet [31, 88, 53], and 28 custom-trained models spanning various architectures [101, 16, 156, 68]. We also include two dynamic models from prior NeuroAI literature—DorsalNet [116] and BLT [158]—both of which have lower brain alignment relative to most dynamic models in our collection (yellow bars in Fig. 2b). This supports the strength and diversity of our model collection.

#### Activation extraction from video frames

To emulate humans watching videos, we need to process the video frames using deep learning models in a specific manner. We refer to it as the inferencing process. One ideal inference would be to enforce *causality*, meaning that the activation *A*_*t*_ at time *t* should not rely on the visual input *I* in the future, *e*.*g*., *I*_*t*+1_ at time *t* + 1. Therefore, the extraction of *A*_*t*_ incurs the processing of contextual visual inputs *I*_*t*−*c,t*_ from time *t* −*c* to time *t* (*c* is the duration of the model’s temporal context and could be infinity), which is extremely costly for non-recurrent models. On the other hand, one can loosen the constraint on causality and allow the model to produce activations that depend both on the past and the future. Concretely, one specific inferencing method is:

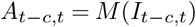

where *M* is the model. In this way, the model processes a whole clip of videos and produces a temporal series of activations *A*_*t*−*c,t*_. Then, *A*_*t*−*c,t*_ is assumed to be evenly distributed across the timestamps to obtain the activation per timestamp. The next inferencing process will run on *I*_*t,t*+*c*_. The process (what we call “block inferencing”) is far less computationally costly than the aforementioned process (what we call “causal inferencing”), while the two give strongly correlated brain alignment scores (see Fig. S28). Therefore, we adopt “block” inferencing for models. Furthermore, since it is also computationally infeasible to process the videos of minutes at once, we limit the context duration to *c* =4 seconds if the model allows flexible input duration. Otherwise, we set the context duration to the model’s fixed input duration. Finally, each model processes the video frames with the same FPS when it was pre-trained, regardless of the FPS of the videos themselves. For static image models, we fixed the processing FPS to be 25. When a video model processes an image, the image is treated as a static video for 1 second.

#### Recording and downsampling layer activations

We record the model activations from all layers for the subsequent analyses, which consume high disk storage. Therefore, we downsample activations from individual layers to a feature vector of fixed length 5000 (if the layer activation had more than 5000 values) using sparse random projections [94]. This projection has been shown to give highly correlated alignment scores with those with the original activations [57]. This also reduces the variance induced by different input sizes when performing the regression analysis.

### B.3. Neural alignment computation

#### Temporal mapping between model activations and fMRI recordings

We map the model activations to the fMRI recordings using a fixed linear transformation over time. This procedure embeds the assumption that the mapping between models and the brain should be invariant across time and datasets, thus making the results more generalisable. At the same time, it keeps the computation cost and overfitting effect at the minimal level. For learning this linear transformation, we need to temporally align the model activations and fMRI signals. As mentioned above, the model activations are generated per frame, with the same sampling FPS as when the model was originally pretrained. On the other hand, all of the fMRI datasets we used have 1 volume per second. To align the model activations to the same temporal resolution as the fMRI has, we average the activations with the 1-second time windows. Furthermore, to simulate the fMRI sampling process, we convolute a hemodynamic function implemented by *nilearn* (originally from SPM, with 5 seconds as the peaking time parameter) to the timeseries of model activations. This method produces better fMRI alignment in our preliminary small-scale tests (see Fig. S29).

#### Cross-validated regression for brain alignment

The brain alignment is defined using correlation over time between the model predictions and the normalised fMRI signals. The predictions are generated from regression, and the whole process is cross-validated for estimating variance. As shown in Fig. 2a, the videos from all datasets are stacked along the time dimension. We segment this long sequence into 15-second video clips and split them into training, testing, and validation sets (80%, 10%, 10%). RidgeCV regression [128] is used to regress the activations from each layer individually to all voxels in the training set. We use 9 different regularisation weights ranging from 10^−4^ to 10^4^ (evenly distributed on a log scale) for the RidgeCV, which picked the best parameter through an approximate leave-one-out error estimation. Then, the regressions were evaluated on the validation set. For each voxel, the best aligned layer (with the least regression error) is determined by the validation set. Finally, the alignment of the model is evaluated as the correlation between model predictions and the ground-truth fMRI signals on the testing set, with the predetermined voxel-layer correspondence from the validation set. The whole process is repeated 10 times for each model with Monte Carlo sampling on the splits to estimate the mean and variance of brain alignment. The choice of using a 15-second clip duration gives a high cross-dataset consistency (0.781) of the motion index map (Fig. 4f). A longer clip duration leads to a substantial drop in consistency (Fig. S23), but still gives qualitatively similar results (SI C). Finally, these brain alignment scores are all normalised by the voxel-level internal consistency (divided by the ceiling) described in Methods B.1.

#### Layer selection and estimation of model-mapped hierarchy

We hypothesise that the hierarchical layer mapping from the models reflects the true computational hierarchy in the brain. For each model, we only subsample at most 15 layers for brain alignment for the sake of saving storage and runtime. For models with block modules, we try to include all the blocks. Otherwise, we evenly sample layers from the whole model. As mentioned above, there is an assigned layer for each voxel from the regression procedure. We compute the relative position of this layer as *k/L*, where *k* is the layer position and *L* is the total number of layers considered for this specific model. Despite prior works claiming that the layer correspondence is inaccurate [153, 125], we get a rather consistent layer assignment by averaging the 20 most brain-aligned dynamic models (split-half consistency over visually driven voxels: 0.839±0.022). This relative layer assignment is used for analyses in Fig. 2e (Fig. S6), Fig. 4e. To compare the model-mapped hierarchy with an existing reference, we used the region connectivity suggested by *Rolls et al*. [142]. We applied an annealing-based ordering procedure 100 times, recording the position of each region in the hierarchy on each run, and then averaged these positions to obtain a probabilistic hierarchy estimate for each region. We then correlated these reference hierarchy values with the model-mapped hierarchy values in Fig. 2e.

### B.4. Behavioural alignment computation

#### Behavioural decoders

To compare behavioural judgements on action recognition videos between humans and models, we need to enable models to produce judgements. We achieve this by training behavioural decoders using data distinct from the human behavioural experiments. Specifically, for each presentation condition (RGB and biological motion), we collect 100 samples per action class, resulting in 1,000 samples per condition for fitting the behavioural decoders. The two decoders are applied to the corresponding groups: one for normal and shuffled RGB videos, and the other for normal and shuffled biological motion videos. For each model, the layer that achieved the highest testing accuracy on the fitting data is selected to generate judgements for the stimuli used in the human behavioural experiments. Both model and human judgements are compared to the ground truth labels, and their accuracies are compared in SI A.7.

#### Human error pattern alignment

In addition to comparing accuracy between models and humans, we are more interested in whether they exhibited similar error patterns. To analyse this, we apply the one-versus-other object-level performance metric (*O*_2_) [134], adapted to our context as an *action-level* metric. The *O*_2_ matrix is defined as:

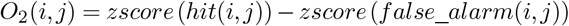

where *i, j* represent indices of the *action classes* for the ground truth and model/human choice, and *zscore* is the inverse of the cumulative Gaussian distribution. The hit rate (*hit*) is the proportion of trials where videos of action *i* are correctly labelled when presented against distractor action *j*, and the false alarm rate (*false*_*alarm*) is the proportion of trials where videos of action *j* are incorrectly labelled as action *i*. Effectively, the *O*_2_ matrix encodes the sensitivity measured by *d*′ for each video-choice combination. The model-human alignment is then computed as the correlation between elements from the model *O*_2_ matrix and the human *O*_2_ matrix, excluding the *i* = *j* entries (as we are primarily interested in the error patterns rather than correct classifications).

### B.5. Decoding performance on cognitive tasks

#### Cognitive task datasets

We consider 10 tasks as the hypothesised functional objectives that drive the visual brain during natural dynamic vision. These tasks and their corresponding datasets are: *object recognition* (`imagenet` [88]): classifying the object categories; *face recognition* (`vggfacev2` [18]): recognising the faces of famous people; *pose estimation* (`majajhong2015` [70]; here we only use the pose data, without the neural recordings): assessing the position and rotation of objects; *social understanding* (`mcmahon-social` [111]): classifying whether two people are interacting/communicating; *appearance-focused action recognition* (`kinetics400` [79]): general-purpose action recognition that includes action classes that are easy to classify based on objects, *e*.*g*., playing ping-pong; *motion-focused action recognition* `smthsmthv2` [62]): action recognition that focuses on solely the movement, *e*.*g*., putting [something] on [something] (the [something] can be arbitrary objects); *egomotion estimation* (`airsim` [116]): assessing the translational and rotational speeds of objects; *direction discrimination* (`ding2012` [33]): assessing the moving direction of a group of dots with randomness; *motion-only action recognition* (`afd` [75]): recognising the action from optical-flow-embedded white-noise stimuli; *biological-motion-based action recognition* (`hdm05` [121]): recognising the action from moving joints confounded by a background of static random dots. For `hdm05`, we project the 3D joints onto rotations of -45, 0, and 45 degrees and add random dots of the same size to the background. This makes it impossible to recognise the positions of joints from a single frame. The same property also applies to `afd` and `ding2012`, making them purely dynamic. The purely static datasets include `imagenet`, `vggfacev2`, and `majajhong2015`, all including only image stimuli. The rest of the datasets have dynamic stimuli, but studies have been arguing that solving these tasks largely involves processing static information [75, 40]. Fig. S10 shows the samples for every task and how these tasks span the spectrum from purely static to purely dynamic information processing (SI B.1). This spectrum of tasks therefore provides a good opportunity to study the interplay between the two kinds of dichotomous information. All sample images shown in Fig. S10 and Fig. 3 are drawn from the original datasets described above. For any images containing humans, we regenerated the visuals using ChatGPT [127] to ensure anonymity.

#### Subsampling of the datasets

We have to perform subsampling on the task datasets to limit computation and storage costs. For each regression task, including `airsim` and `majajhong2015`, we randomly sample 5000 instances from the dataset. For other classification tasks, we cap the number of classes to be 100 and the number of samples per class to be 50.

#### Cross-validated decoding performance of tasks

Even if the models are not originally optimised for these cognitive tasks, we can measure how much their activations *support* them, similarly to neural decoding. This is done by learning regression/classification from the model activations to the various task labels, either continuous or categorical. We apply a cross-validation process that is identical to that of evaluating brain alignment, with the proportions of training, testing, and validation sets to be 80%, 10%, 10%. The layer selection for decoding is again done with the validation set. However, a major difference is that when selecting the layers, we only consider the top layers from the models. The representation at the top layers is constructed progressively throughout the whole model due to their hierarchical structures. In other words, the task supported by the top layers shapes the representation of all preceding layers. Therefore, we choose to consider the top 20% layers (out of the subsampled layers) and take at least 2 top layers from each model.

### B.6. Task relevance computation

#### Task relevance

The relevance of a single task to the brain (or brain regions) is simply defined as the explained variance (or R-squared) from correlating the task performance and the whole-brain (or regional) alignment scores across all tested models. A high relevance indicates that by increasing the performance of models on this task, we expect their neural or behavioural alignment to also increase. This further suggests that this task might serve as a functional objective for the computation across the brain or in those specific regions. Similarly, for behavioural alignment, a high relevance implies that the hypothesised cognitive task may drive the model’s alignment with human behavioural outputs. It is worth noting that sometimes the task performance can be negatively correlated with the brain alignment. We regarded these cases as artefacts from the limited size of our model pools. Therefore, we constrain all the regressions used for task relevance computation to have non-negative weights.

#### Joint task relevance

A single area in the brain might serve for multiple different functional goals achieved in the downstream areas. To consider the relevance of multiple tasks in combination, we regress the performance on these tasks to the brain or regional alignment. Then, the joint relevance is defined as the adjusted explained variance (or adjusted R-squared) of this regression:

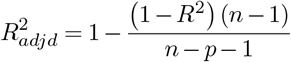

where *R*^2^ is the explained variance (or R-squared). The adjustment takes into account the number of regressors used. Analogically, the joint relevance reflects how accurately the brain alignment can be predicted from the performance of given tasks, therefore how these tasks drive the neural processing in combination.

#### Residual task relevance

Since the underlying mechanism of solving two tasks can be similar, *e*.*g*., face and object recognition both involve integrating shape and texture information across space, the relevance of the two tasks might overlap with each other. To remove the contribution of a specific task in the joint task relevance, we considered the unique explained variance of the remaining task. Generally, if the whole set of tasks considered is *T* and the set of tasks we want to remove is 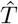, the residual task relevance of 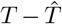 is 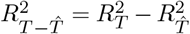. If the joint relevance of *T* can be fully explained by the subset 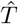, the residual task relevance 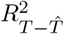 should be close to 0. For verifying the statistical significance of the residual task relevance, we conduct permutation tests with 1000 resamples.

#### Eigenanalysis of Voxel-Level Task Relevance

In Fig. 3g, we visualise the 2D eigenspace of visually driven voxels across the cortex, based on their task relevance profiles. Specifically, each voxel is represented as a point in an *M* -dimensional space, where *M* = 10 corresponds to the number of cognitive tasks, and the coordinates are given by the relevance for each task, *i*.*e*., the vector 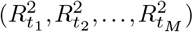. Applying eigendecomposition to the voxel-by-task relevance matrix, we find that the first two principal components together explain 91% of the total variance, with a balanced contribution from each component (47% and 44%, respectively), indicating that the space is evenly spanned in two dimensions. Furthermore, projecting each task vector onto this eigenspace reveals two dominant, approximately orthogonal axes that correspond to object-related and motion-related processing. These axes capture the principal semantic structure underlying the voxel-task relationships.

#### Motion index of information processing

We consider an intuitive measure of how a brain region processes static versus dynamic information: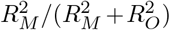, where *M* represents `afd` and *O* represents `imagenet`. This measure ranges from 0 to 1, indicating from processing only static information to processing only motion information. At the level of 0.5, it indicates an equal processing of both sources of information. To make it more intuitive, we map the range from 0% (purely static) to 50% (equal) and then back to 100% (purely dynamic) in Fig. 4f and Fig. 5f.

## C. Quantification and statistical analysis

All analyses are conducted in Python using two-tailed statistical tests. For paired samples (*e*.*g*., individual model scores), Student’s t-tests are applied, while for non-paired samples (*e*.*g*., model group scores and task relevance), Mann-Whitney U tests are used. Multiple comparisons across tasks, voxels, or regions are corrected using the false discovery rate (FDR) method. To estimate the 95% confidence intervals for task relevance, bootstrapping with 1,000 resamples is performed over model scores. On the other hand, the confidence intervals of model scores are directly based on estimated standard deviations from different data splits. To assess whether task relevance or remaining task relevance is significantly above zero, permutation tests with 1,000 resamples are conducted. Statistical significance thresholds are defined as follows: not significant (n.s.), *p>* 0.05; ∗*p<* 0.05; ∗∗ *p<* 0.01; ∗∗ ∗*p<* 0.001, unless stated otherwise.

## Acknowledgments

This work was supported by the Swiss National Science Foundation (SNSF) under Grant *No. 10*.*003*.*772*. We thank Ben Lönnqvist, Johannes Mehrer, Nancy Kanwisher, and members of the EPFL NeuroAI Lab for helpful feedback. We also thank Yitao Xu and Huzheng Yang for valuable discussions.

