## Supplementary materials for "Diverse Perceptual Representations Across Visual Pathways Emerge from A Single Objective"

### Supplementary Information

#### A. Additional details on brain alignment

***A.1. fMRI datasets.*** Table 1 summarises the statistics of the fMRI datasets used in this study. For most datasets, the number of repetitions per stimulus is limited to one. Ideally, per-subject brain alignment would be more informative, as group-level averaging can obscure fine-grained spatial information across individuals. However, due to the low number of repetitions, we chose to average across subjects to obtain more robust signal estimates and conducted group-level analyses.
Three of the five datasets used in this study are audio-visual, providing reliable signals in auditory regions as well. However, since this paper focuses exclusively on visual modelling, even the best-performing models exhibit relatively low alignment in the vicinity of the auditory cortices (Fig. 2c).

***A.2. Alignment of untrained models.*** To show the necessity of task optimisation, we selected the top models from each model group and compared their brain alignment to that of their untrained counterparts (Fig. S1). The untrained versions were obtained by randomly reinitialising the model weights.

***A.3. Alignment of all models.*** Figure S2 presents detailed neural alignment scores across the visual system for individual models, alongside their parameter counts and processing speeds (FPS). It is evident that neither model size nor processing speed correlates with alignment performance. Notably, some older architectures, such as AVID-Audioset and I3D-Nonlocal, achieve alignment scores comparable to more recent models like V-JEPA.

***A.4. Ventral stream alignment in both dynamic and static datasets.*** To assess model alignment with the ventral visual stream under static image viewing, we evaluate models on macaque electrophysiological recordings from areas V1, V2, V4, and IT [177, 107]. In these experiments, macaques were trained to fixate at the screen center while images were presented, and neural responses were averaged over time to obtain mean firing rates for each stimulus. For V1 and V2, 450 naturalistic and noise images were shown, with recordings from 102 (V1) and 103 (V2) electrodes [177]. For V4 and IT, 5,760 naturalistic images were presented, with recordings from 192 (V4) and 480 (IT) electrodes [107]. Each stimulus was repeated at least 20 times to ensure reliable neural signals. Further methodological details are available in the original publications.
Despite their focus on temporal processing, dynamic models still perform on par with static models in ventral stream regions (Fig. S3c). Image recognition models have long been regarded as state-of-the-art for predicting primate ventral stream responses during static image viewing [147, 148] and, while they remain superior under static conditions (electrophysiological data [147, 107, 177]), they do not perform as well as action recognition and text-video models in predicting dynamic video-induced brain activity. When averaged across all conditions, alignment scores for image and dynamic model families become statistically indistinguishable.

***A.5. The best dynamic model vs. the best static model.*** Figure S4 illustrates the alignment of V-JEPA, the best-performing model on the dynamic datasets, to both hemispheres and medial views, complementing Figure 2c. It exhibits prominently high alignment in dorsal regions near MT, V3A, and V6, as well as decent alignment in higher-level ventral areas, the SPL, and regions near the pSTS. Alignment is lower in the IPL and further decreases in more anterior portions of the STS and areas associated with auditory processing. In contrast, the top model for the static ventral stream datasets, ConvNeXt-L, exhibits notably lower alignment, particularly in dorsal regions.

***A.6. Details on model-mapped hierarchy.*** Figure S6 shows the complete hierarchy map across both hemispheres. The hierarchical progression along all visual streams is clearly delineated: the ventral stream culminates near the PPA, the dorso-dorsal stream extends to the SPL and areas near the postcentral sulcus, and the ventro-dorsal stream ascends toward regions around the pSTS, with slightly higher hierarchical levels observed in the right hemisphere.
The most brain-aligned layers are typically in the latter half of the model, which might suggest that processing in earlier layers relates more to retinal and subcortical processing.
Figure S7 shows a weak but significant correlation between the model-mapped hierarchy and neural alignment scores. Notably, lower-alignment voxels tend to be best predicted by higher model layers. However, the low correlation coefficient indicates that this effect is modest. Combined with the hierarchical patterns observed across the visual streams, we argue that the model-mapped hierarchy nonetheless accurately reflects the increasing computational complexity along the visual hierarchy.

***A.7. Behavioural alignment in terms of recognition accuracy.*** Figure S8 presents action recognition accuracy across various video conditions. Many dynamic models achieve human-level performance, whereas static models generally underperform. Importantly, the pattern of accuracy changes across conditions in dynamic models closely mirrors that observed in humans. This model-human alignment complements the error pattern alignment shown in Figure 2g.

### B. Additional details on task relevance

***B.1. Cognitive tasks.*** The full list of cognitive tasks and their sample stimuli is shown in Fig. S10. Model performance across these tasks exhibits varying degrees of correlation, as illustrated in the similarity matrix in Fig. S11. This matrix reveals the extent to which each task leans toward static or dynamic processing. Importantly, these task scores span a higher-dimensional space, with the top four principal components explaining 52%, 26%, 14%, and 3% of the variance, respectively. This indicates a richer structure than the largely two-dimensional task-relevance space shown in Fig. 3g. This demonstrates that the two-dimensional task-relevance space is non-trivial and meaningfully captures the processing nature of the brain.

***B.2. Other combinations of object and appearance-free motion recognition.*** In the main text, we presented results from combining object recognition (`imagenet`) with motion-only action recognition (`afd`), which yielded the highest joint task relevance.

To determine which form of motion information contributes, we further test an additional high-level appearance-free motion recognition task (biological-motion-based action recognition) and a low-level task (direction discrimination). Combining the former with object recognition still explains substantial variance ($adjd.\ R^2 = 61\%$, Fig.3c; $adjd.\ R^2 = 38\%$, Fig. 3d) and explains away relevance across tasks (Fig.3e). In contrast, combining low-level motion recognition explains substantially less variance ($adjd.\ R^2 = 42\%$, Fig. 3c), suggesting that recognising high-level dynamics is critical for this joint processing.

Figure S12 shows several other task combinations with statistically similar relevance to the combination of `imagenet` and `afd`. Crucially, these effective combinations consistently include both high-level object and high-level motion recognition capacities. For example, `vggfacev2` (purely static) paired with `afd` (purely dynamic), or `kinetics400` (primarily static with some dynamic components) paired with `airsim` (emphasising dynamic information). These findings further highlight the nature of object-motion joint processing in the brain.

***B.3. Stream ROIs from the Glasser atlas.*** The regions along different visual streams analysed in the paper are defined according to the Glasser atlas (Fig. S14).

***B.4. Task relevance for MT and FFC.*** To demonstrate the contribution of individual tasks to combined task relevance, we analyse the ventral region FFC and the dorsal region MT (Fig. 4B). Motion-only action recognition shows a significantly higher correlation with brain alignment in dorsal MT compared to ventral FFC ($r = 0.70$ vs. $r = 0.43$; $p < 0.01$). In contrast, static object recognition has stronger relevance for ventral VVC over dorsal MT ($r = 0.71$ vs. $r = 0.42$; $p < 0.01$). Overall, combined task relevance is the most predictive, best explaining brain alignment in both regions ($r = 0.91$ for FFC and $r = 0.90$ for MT).

***B.5. Increasing complexity of motion representations along the hierarchy.*** The apparent regional hierarchy in motion processing is indeed reflected in the model's neural mechanisms. Specifically, the layer best corresponding to each region—i.e., the computational depth—strongly correlates with *motion complexity* (Fig. S16; methods detailed in SI B.6). This measure contrasts how relevant low-level motion tasks such as direction discrimination are compared to high-level motion tasks such as action recognition. Analogous to the well-established object processing hierarchy in the visual ventral stream [174, 147], this suggests the existence of a hierarchical motion processing network. Taken together, these findings support the view that object and appearance-free motion recognition are core *representational* principles implemented throughout the visual streams.

***B.6. Motion complexity.*** The motion complexity is defined using the residual task relevance: the motion-only action recognition (`afd`) is a high-level motion processing task, because it requires integrating over a longer time scale and achieving more complex invariance. On the contrary, direction discrimination (`ding2012`) and egomotion estimation (`airsim`) are lower-level. To quantify the extent to which a brain region processes exclusively high-level motion information, we examine the proportion of residual relevance in `afd` after regressing out `ding2012` and `airsim`. In other words, this measure reflects the relevance of high-level motion specifically, relative to the total relevance across all motion types. Specifically, the proportion is measured by $R^2_{M-L}/R^2_M$, where $M$ represents `afd` and $L$ contains `ding2012` and `airsim`.

***B.7. Behavioural relevance of combined recognition.*** The combination of object recognition (`imagenet`) with either motion-only action recognition (`afd`) or biological-motion-based action recognition (`hdm05`) did not reach the relevance level of the highest single task `kinetics400` (Fig. 3d). We hypothesise that certain static features crucial for action recognition, such as body parts and forms, are not fully captured by the object recognition dataset `imagenet`. When substituting `imagenet` with `kinetics400-static`—the same dataset as `kinetics400` but with shuffled frame order—the static-dynamic combinations again achieve relevance on par with or exceeding that of `kinetics400` alone. This finding further supports the explanatory power of these two-task combinations, extending beyond neural alignment to behavioural alignment.

***B.8. Task-based functional localisation.*** The voxel-level task relevance analysis yields functionally localised structures on the cortical surface. Beyond mapping object-motion processing (Fig. 4f) and behavioural correlates (Fig. S17), this approach also enables fine-grained analysis of other functional objectives. As an example, Figure S18 shows the residual relevance of face recognition after regressing out action recognition. The resulting map highlights a series of regions implicated in face processing, including the OFA and FFA.

#### C. Consistency of the results under different temporal generalisation conditions

We chose a temporal granularity of 15 seconds for the train-validation-test splits when establishing neural alignment (Methods B.3). Using longer clip durations implicitly imposes a stronger requirement on the learned mapping to generalise across time. Here, we evaluate whether the main results reported in the text remain consistent under this stricter generalisation requirement.

To test this, we evaluate the case with a clip duration of 60 seconds. Under this condition, dynamic model groups (except for forward prediction models) exhibit a clearer advantage over static models, with action recognition models showing particularly strong performance (Fig. S19). However, the overall alignment levels across all models decrease, as expected, due to the increased difficulty of generalisation under this stricter temporal condition.

The task relevance results exhibit a very similar ordering (Fig. S20) to that shown in Fig. 3c, despite the overall higher values. The combination of object and appearance-free motion recognition continues to account for the relevance of all other tasks, both in combination and individually (Fig. S21). At the regional level, we observe similarly hybrid processing across regions. While the stream-level relevance shows a stronger overall bias toward motion processing under this condition, the relative differences between streams remain consistent (Fig. S22).

Finally, we examine the consistency of the motion index map across different datasets and clip durations. For each clip duration, we split the five fMRI datasets into two halves and conducted motion index analyses on each split. We then computed the correlation between the indices from the two splits to assess split-half consistency. We find that consistency is very high with a 15-second clip duration but decreases to a modest level with 30- and 60-second durations (Fig. S23). However, qualitative inspection of the motion index maps reveals similar spatial patterns even with a 60-second clip duration (Fig. S24), and these patterns remain consistent with the analyses in the main text. A strong robustness of model neural alignment is also observed (Fig. S25).

| Dataset | #participants | #stimuli | Stimulus duration | #repetitions |
|---|---|---|---|---|
| *Berezutskaya et al. 2021* | 30 | 1 (audio-visual) | 6.5 minutes | 1 |
| *Keles et al. 2024* | 24 | 1 (audio-visual) | 8 minutes | 1 |
| *Sava-Segal et al. 2023* | 43 | 4 (audio-visual) | ∼8 minutes | 1 |
| *Lahner et al. 2024* | 10 | 1000 (visual-only) | 3 seconds | 1 |
| *McMahon et al. (2024)* | 4 | 250 (visual-only) | 3 seconds | 2 |

**Table 1. Statistics of fMRI datasets.**

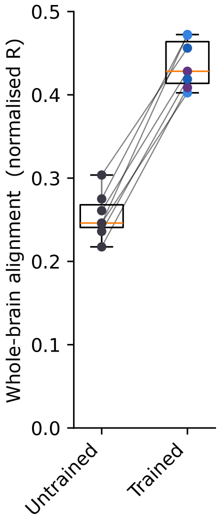


**Figure S1. Comparison between untrained and trained models:** alignment scores of multiple top-performing models from different groups significantly surpass those of their untrained counterparts.

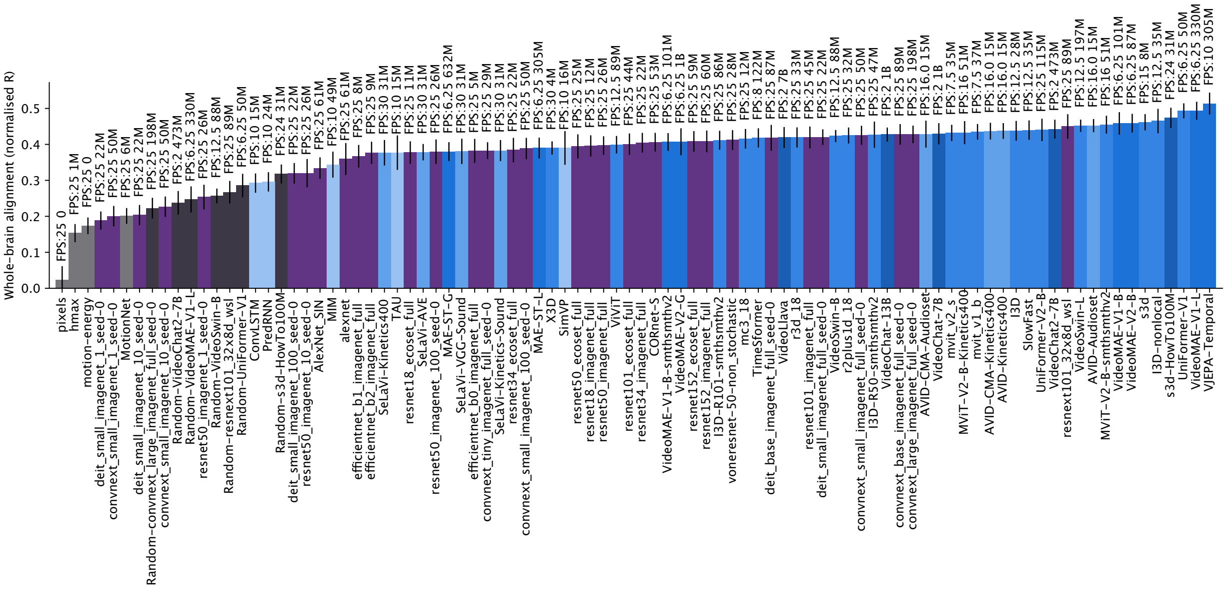


**Figure S2. Whole-brain alignment scores of all models.** The colouring follows Fig. 2b, except that the yellow colours indicate models that we considered but do not belong to any group of interests in this paper. The texts above the bar plots indicate the FPS with which the model was pretrained and the parameter sizes.

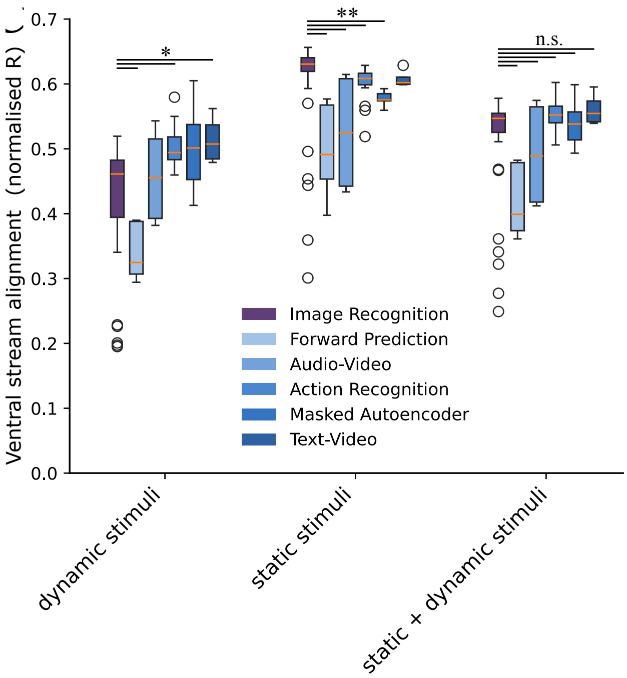


**Figure S3. Ventral stream alignment across stimulus conditions:** the plot shows ventral stream alignment for different model groups on dynamic stimuli (fMRI datasets analysed in this study) and static stimuli (electrophysiological data analysed in Brain-Score [147]), along with their average. While static image recognition models excel in static conditions and dynamic models excel in dynamic conditions, their averaged alignment scores across conditions are on par.

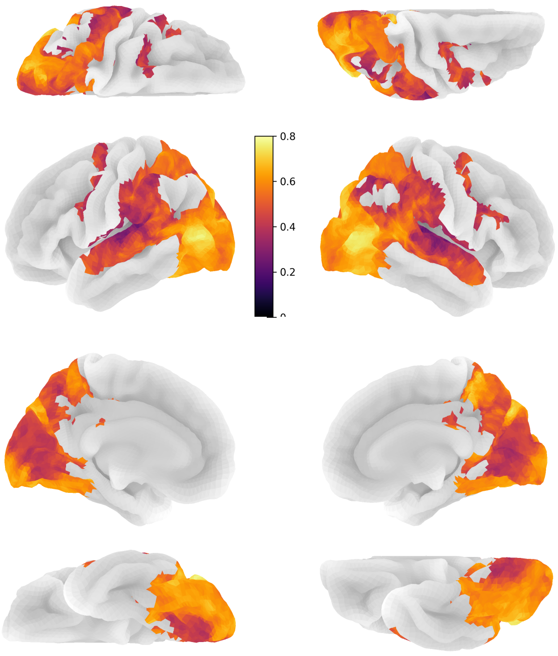

**Figure S4. Normalised brain alignment of V-JEPA:** the data for both hemispheres (partial data in Fig. 2c); from top to bottom: the dorsal, lateral, medial, and ventral views.

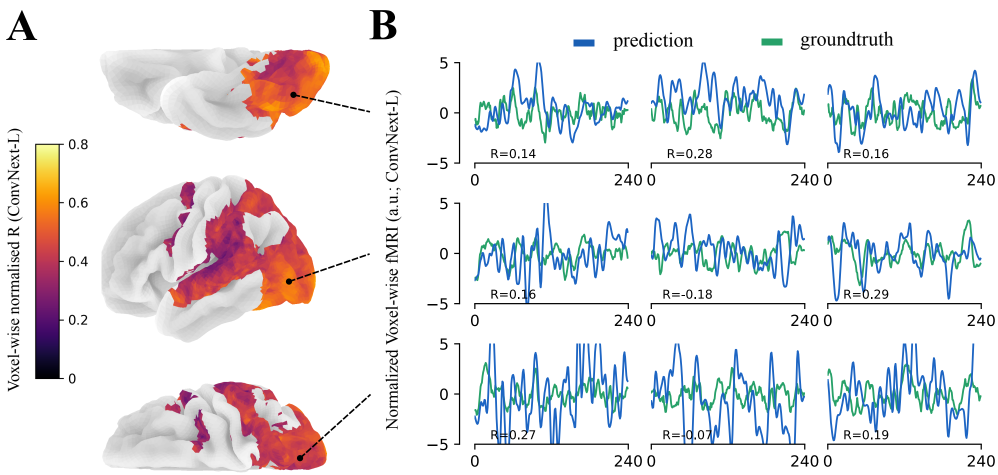


**Figure S5. A. Prediction scores across the voxels of the best static image processing model:** left hemisphere. **B. Predicted signals vs. ground truth of the best static image processing model:** the signals are from three representative voxels on three different movie clips.

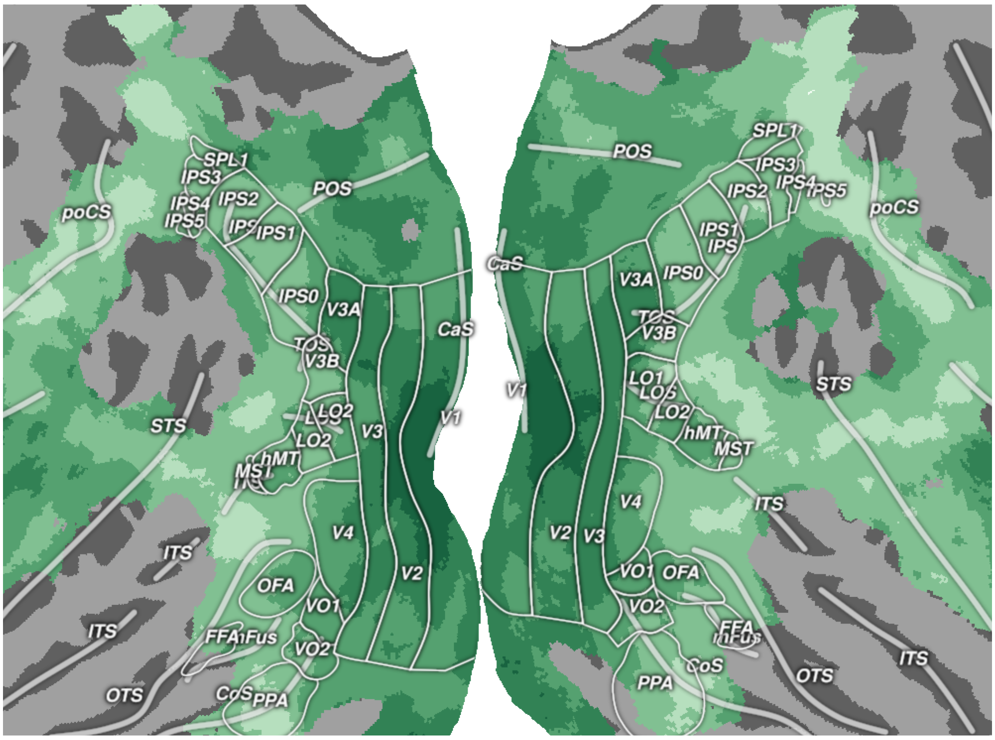


**Figure S6. Model-mapped hierarchy for both hemispheres:** the discretised hierarchy for visually driven regions. The colouring follows Fig. 5e.

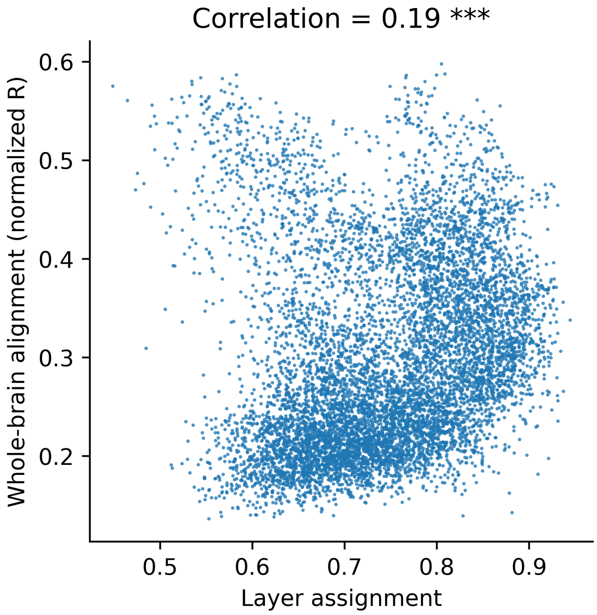


**Figure S7. Model-mapped hierarchy *vs.* alignment scores across visually driven voxels.**

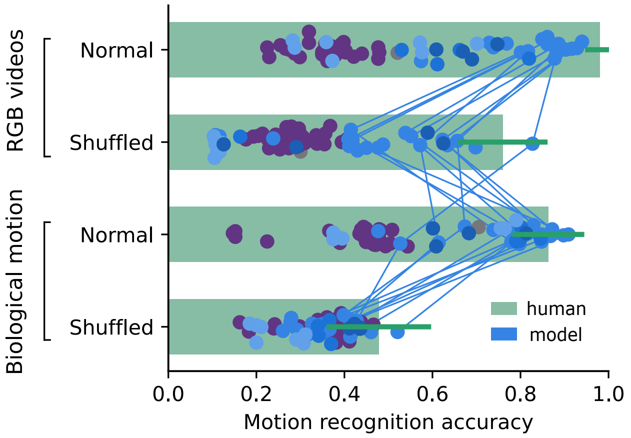


**Figure S8. Human vs. model recognition accuracy:** human performance (green bars) compared to models (blue/purple dots, coloured as in Fig. 2b). Blue lines track the top 10 models across conditions.

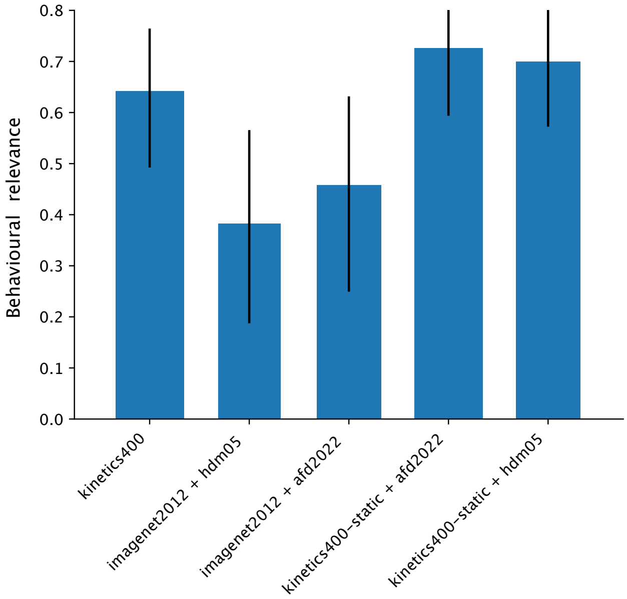


**Figure S9. Behavioural relevance of combined image-based action recognition and appearance-free motion recognition:** compared to Fig. 3d, we add the combinations of `kinetic400-static+afd` and `kinetic400-static+hdm05`, both yielding higher mean alignment than `kinetic400` alone.

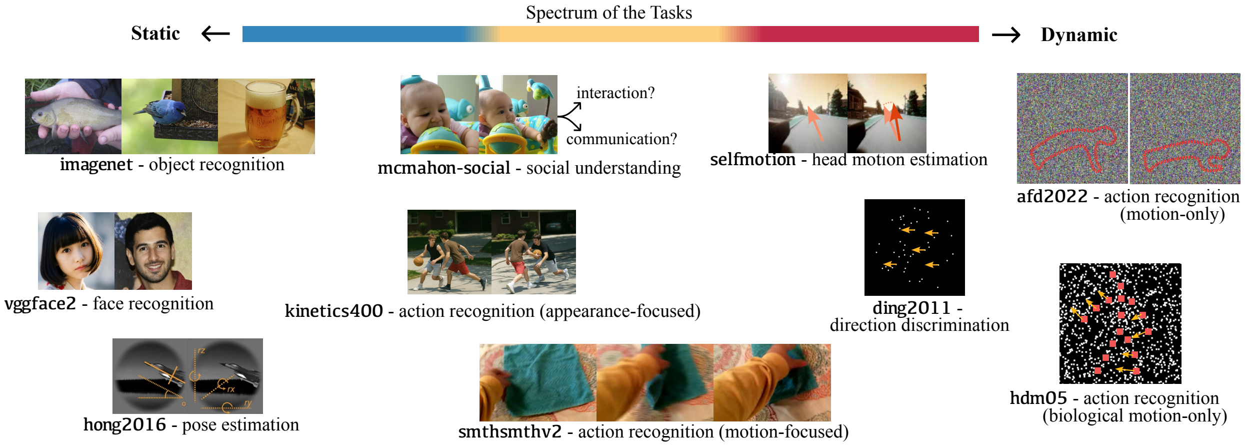


**Figure S10. Samples of all cognitive tasks considered in this paper.**

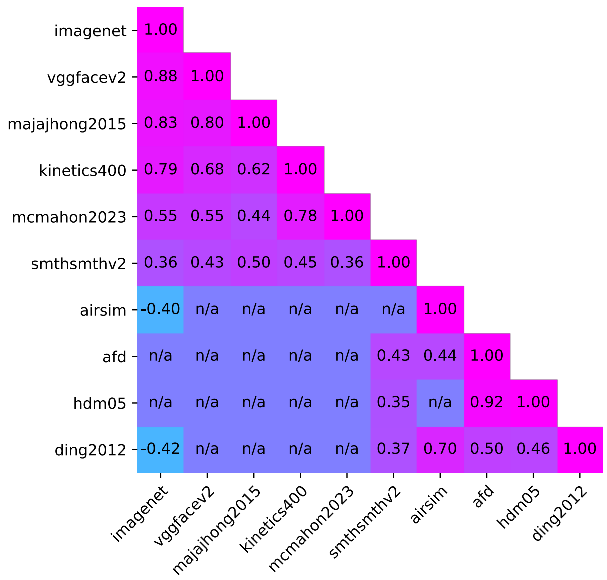


**Figure S11. Correlations between model scores on different tasks:** the “n/a” indicates non-significant correlations.

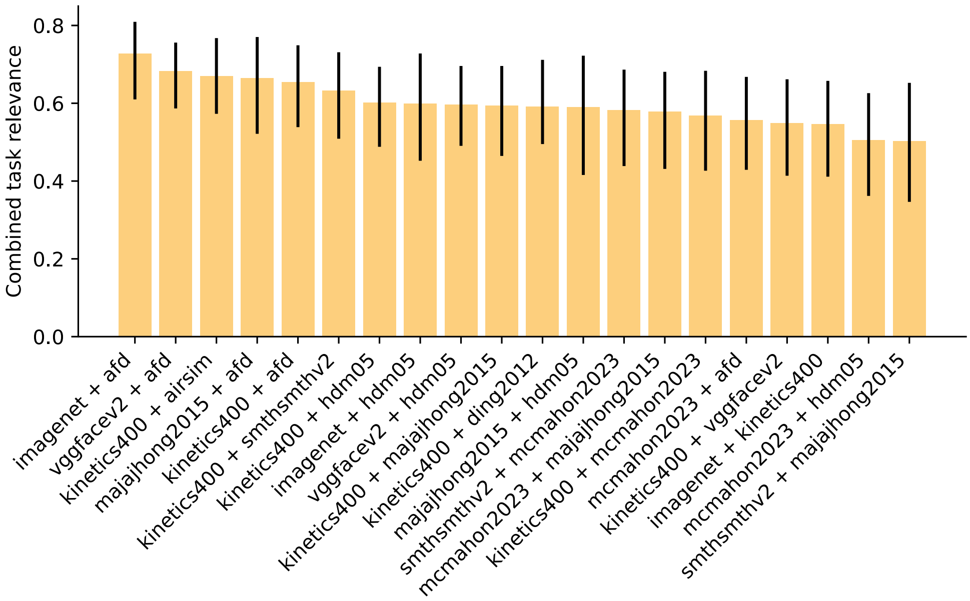


**Figure S12. Task pairs with the highest 20 combined task relevance:** these combinations come from exhaustive research of all possible two-combinations from the set of 10 considered cognitive tasks. Though many of them have similar task relevance to that of the object and motion-only action recognition, the top combinations all share the pattern of jointly processing static object and appearance-free motion information.

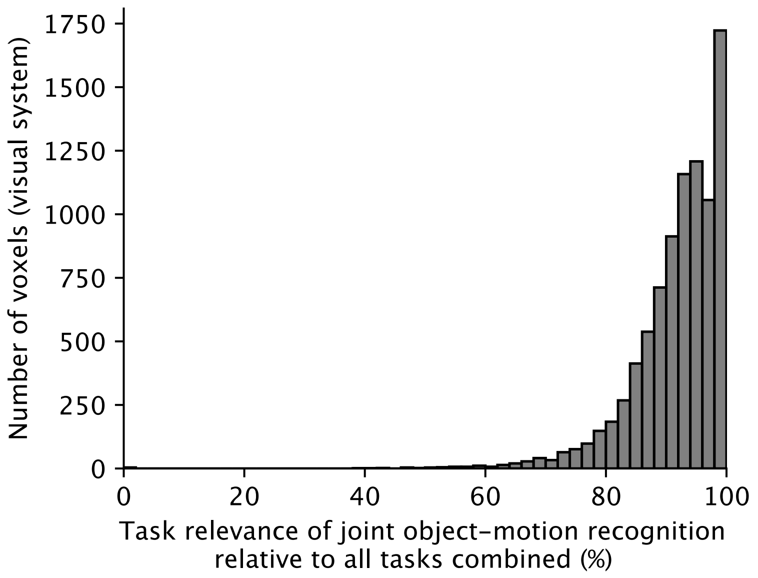


**Figure S13. Voxel-wise fraction of object–motion relevance out of all tasks:** distribution of the proportion of each voxel's task relevance explained by joint object and motion recognition relative to all tasks combined (10 in total; Methods. B.5).

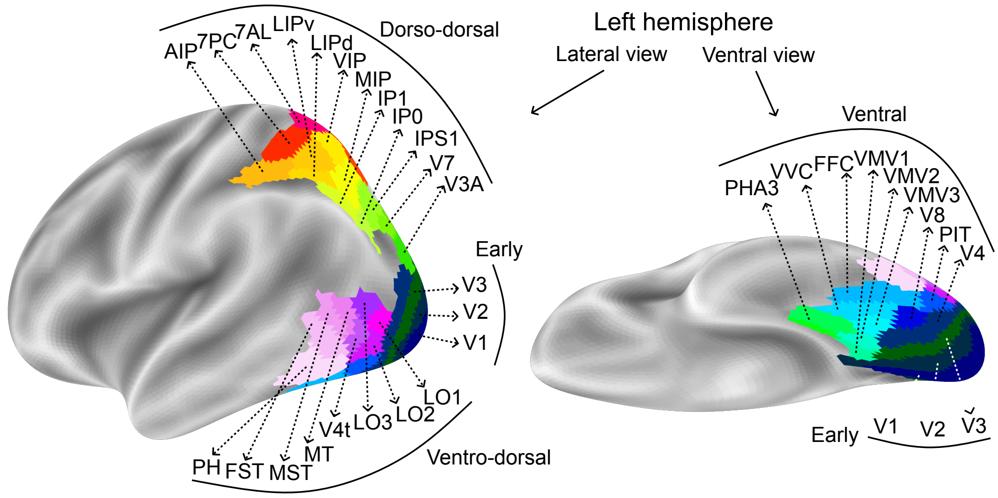


**Figure S14. Stream ROIs from the Glasser atlas:** we conduct the task relevance analysis presented in Fig. 4a at the regional level by selecting regions of interest (ROIs) along the anatomical structures of the ventral and dorsal (including dorso-dorsal and ventro-dorsal) pathways. These ROIs are from the Glasser brain atlas [55].

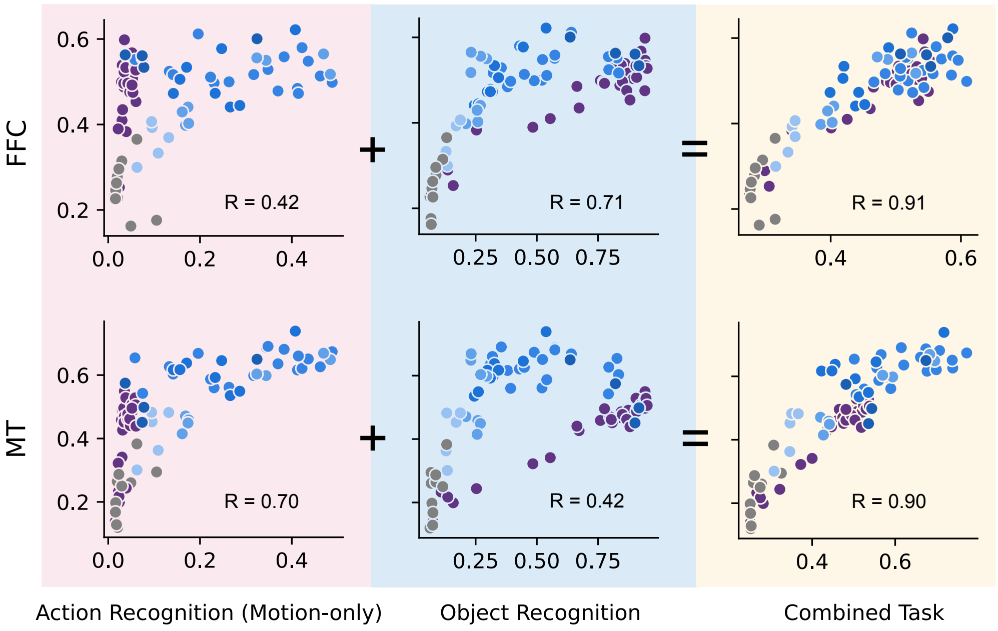


**Figure S15. Individual vs. combined task relevance in FFC and MT:** the correlation plot shows task performance vs. regional alignment for ventral FFC and dorsal MT. Both regions exhibit increased relevance from the combined task, despite their initial bias toward a component task (object recognition or motion-only action recognition).

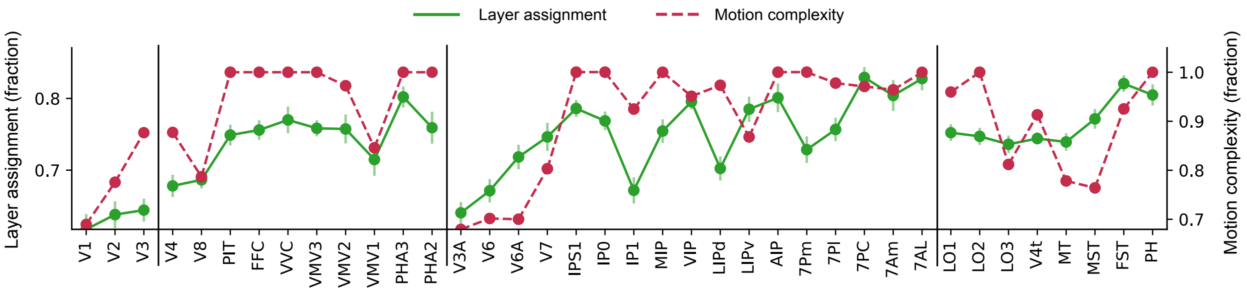


**Figure S16. Individual vs. combined task relevance in FFC and MT:** the correlation plot shows task performance vs. regional alignment for ventral FFC and dorsal MT. Both regions exhibit increased relevance from the combined task, despite their initial bias toward a component task (object recognition or motion-only action recognition).

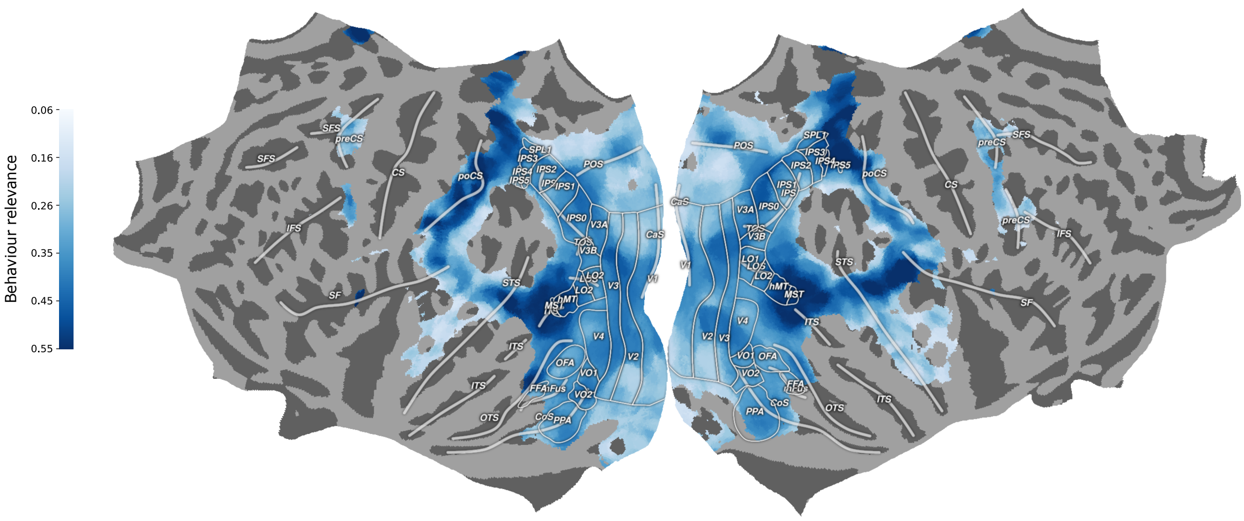


**Figure S17. Brain-wide behavioural relevance map:** the corresponding full map of the partial map shown in Fig. 5d. A slight right-lateralisation is observed.

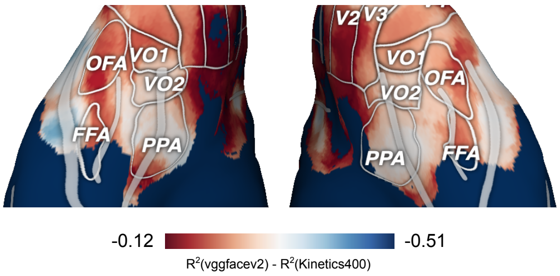


**Figure S18. Computational identification of face processing pathways.** The figure directly contrasts the voxel-wise relevance of *vggfacev2* (face recognition) and *kinetics400* (appearance-based action recognition). The red areas indicate where *vggfacev2* has relatively higher relevance (though still below *kinetics400*), including some known face-selective regions like FFA and OFA.

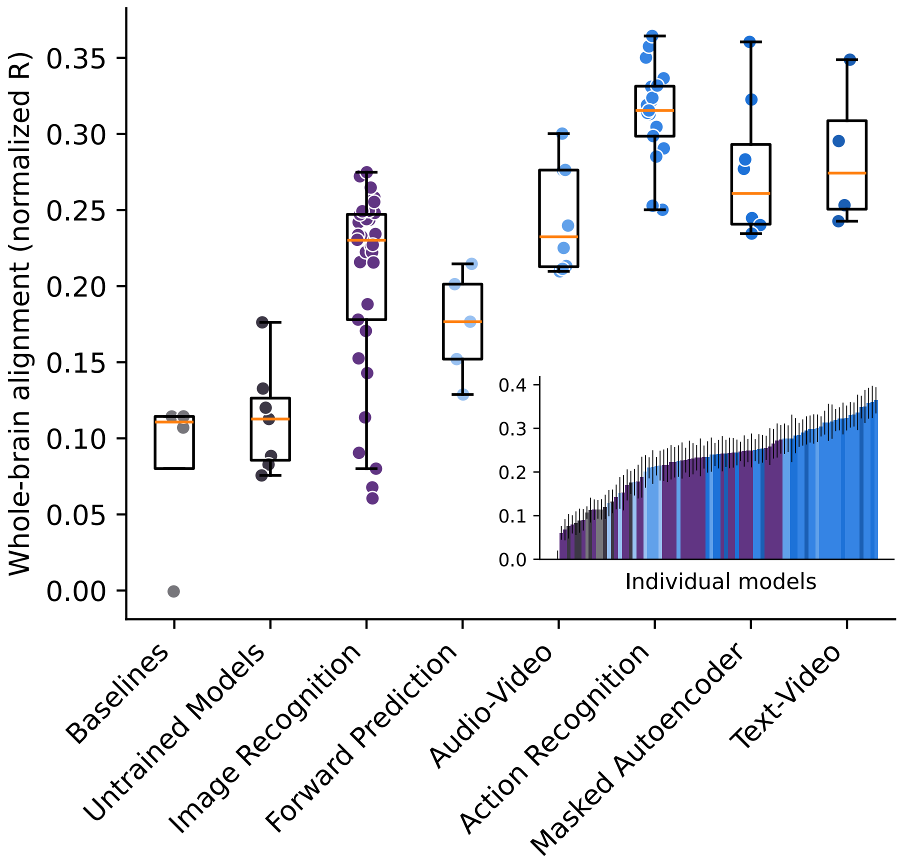


**Figure S19. Comparison of whole-brain alignment of different model groups with 60-second clip duration.** Many temporal model groups show higher alignment than the static image recognition group.

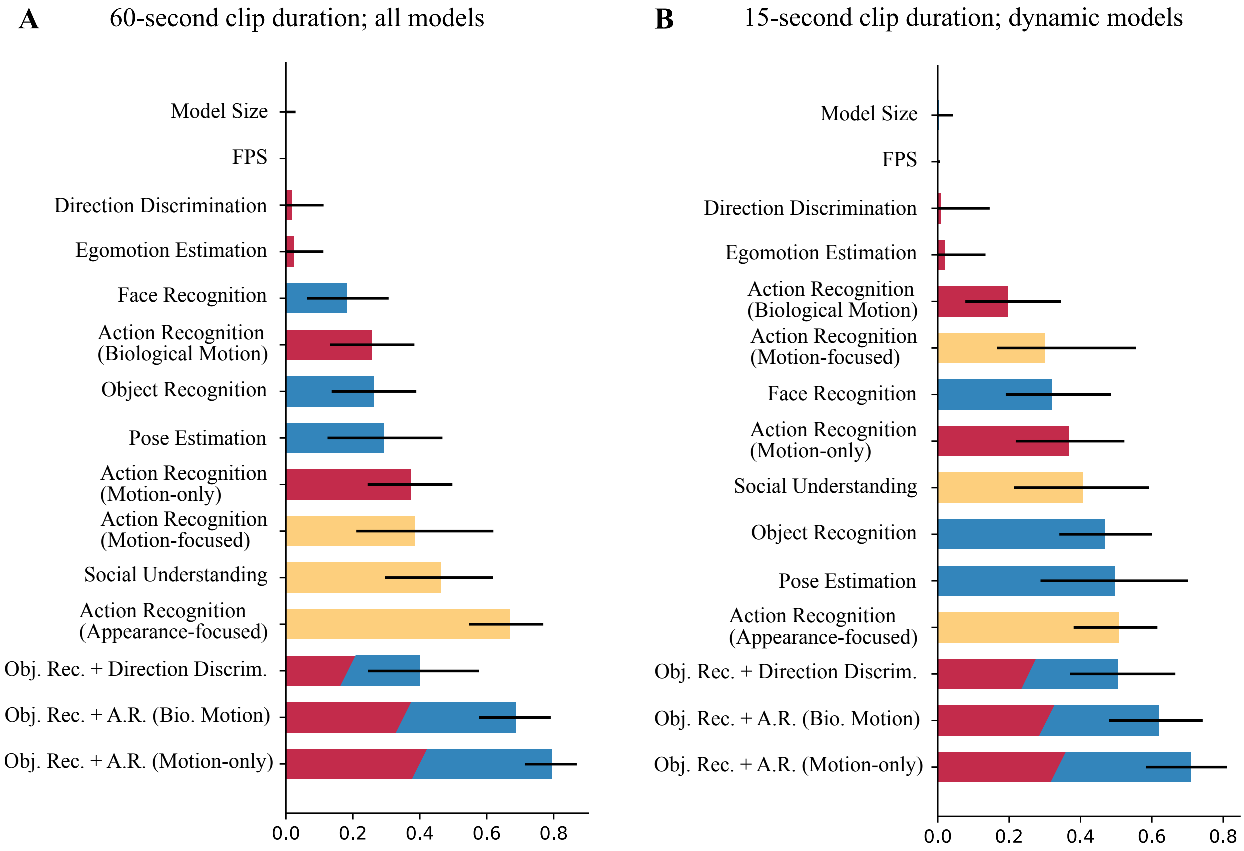


**Figure S20. A. Task relevance with 60-second clip duration and all models. B. Task relevance with 15-second clip duration but only dynamic models.** These qualitative results with 15-second clip duration and all models (default throughout the main text) still hold: 1. A mixed-static-dynamic task achieved the highest relevance; 2. The combined static object and dynamic motion-only action recognition task is significantly more relevant to the brain's processing compared to any individual task.

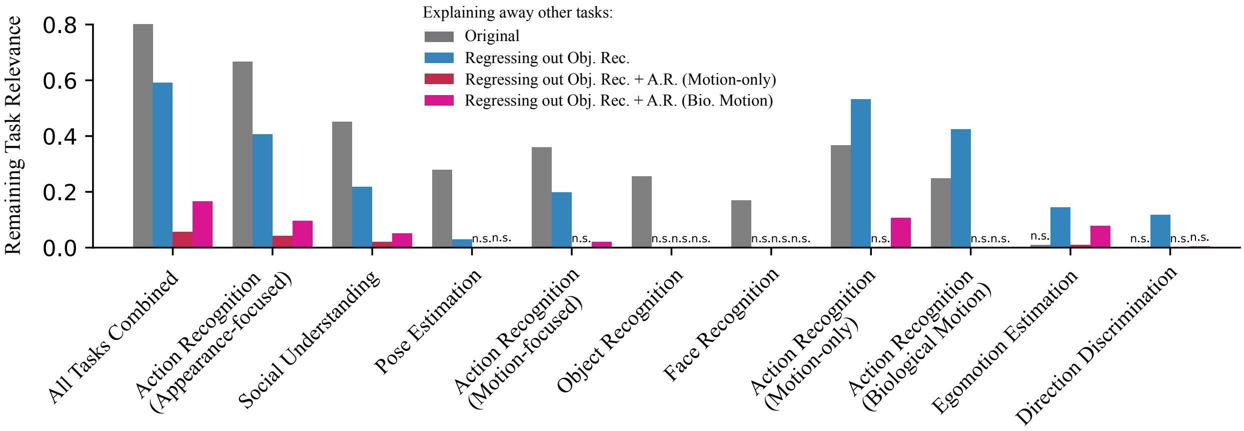


**Figure S21. Residual task relevance with 60-second clip duration.**

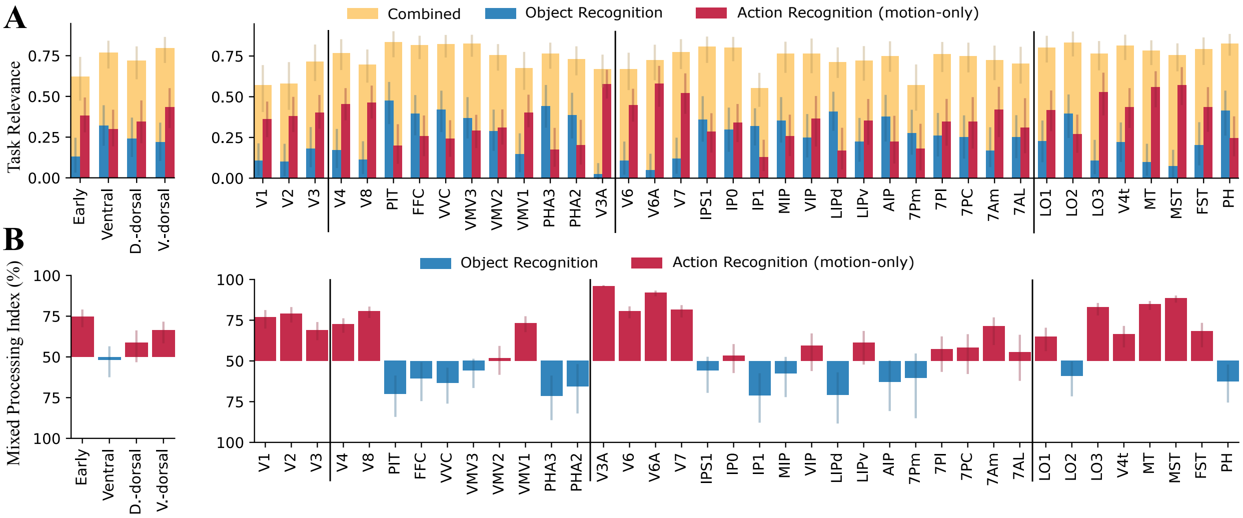


**Figure S22. A. Task relevance of the combined task *vs.* that of the individual tasks for ROIs with 60-second clip duration. B. Mixed processing index of ROIs with 60-second clip duration.**

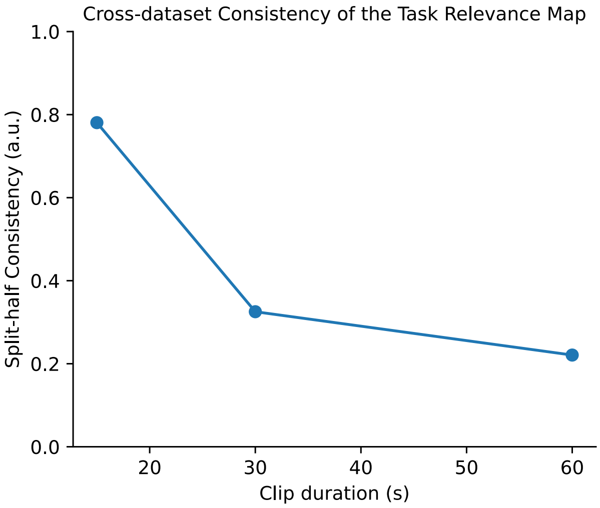


**Figure S23. Split-half consistency of the motion index map across different clip durations:** though the consistency drops substantially with 30/60-second clip durations, the motion index maps are qualitatively consistent, as shown in Fig. S24.

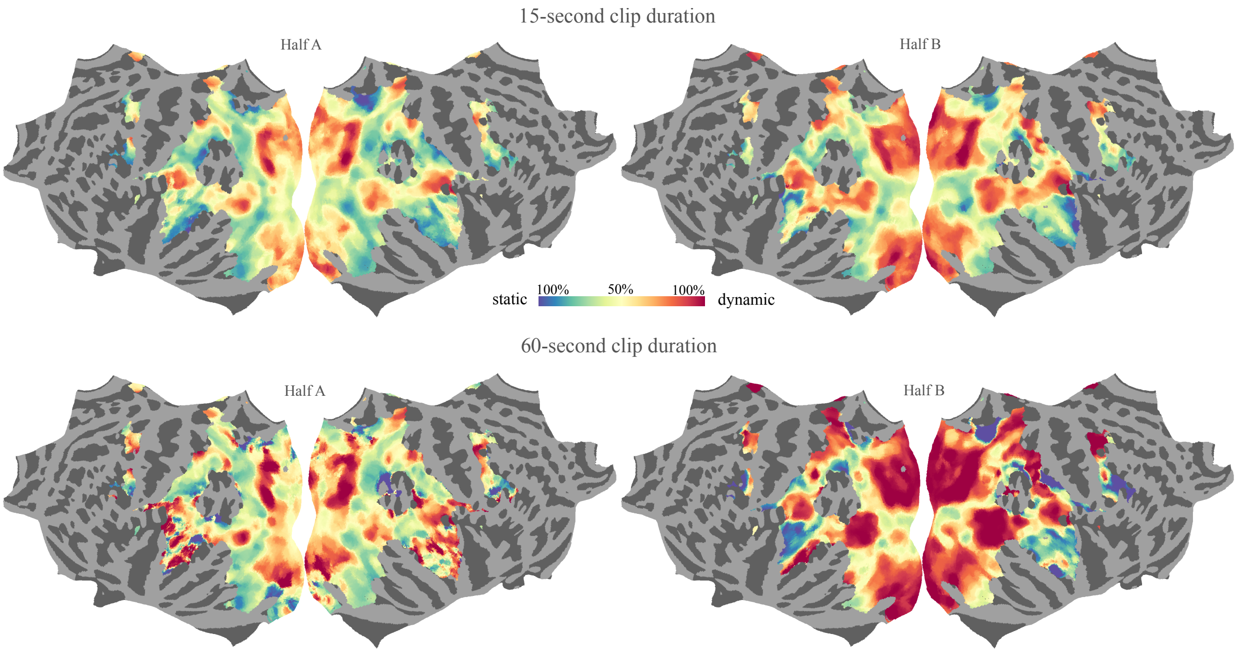


**Figure S24. Examples of task relevance maps generated from two mutually exclusive halves of the datasets:** one single relevance map is generated solely from one half of the datasets. The top and bottom rows, respectively, show the results from using 15-second and 60-second clip durations. All halves give qualitatively similar results on static-dynamic networks, with the 60-second ones indicating stronger dynamic processing.

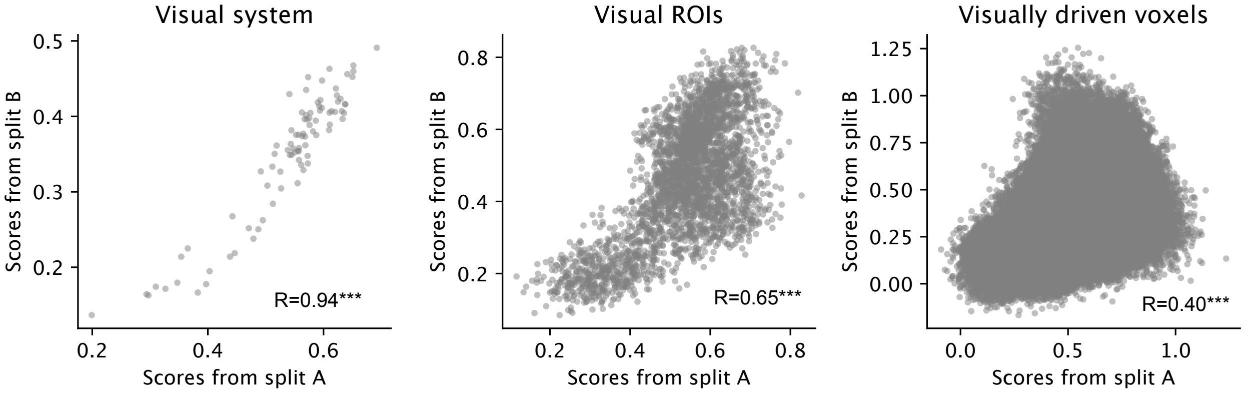


**Figure S25. Neural alignment scores across models on two mutually exclusive halves of the datasets:** scores on the two splits exhibit significant correlations across different levels of regional coarseness.

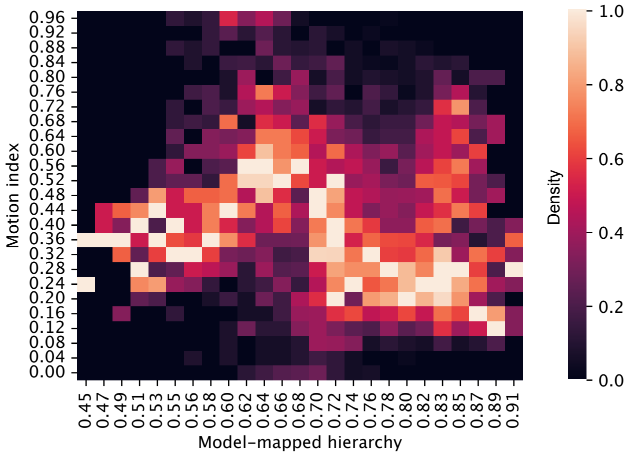


**Figure S26. Unfiltered layer-wise distribution of motion index:** Raw layer-wise distributions of the motion index without filtering. Among the 25 binned levels of the hierarchy, unimodality holds in all but one case.

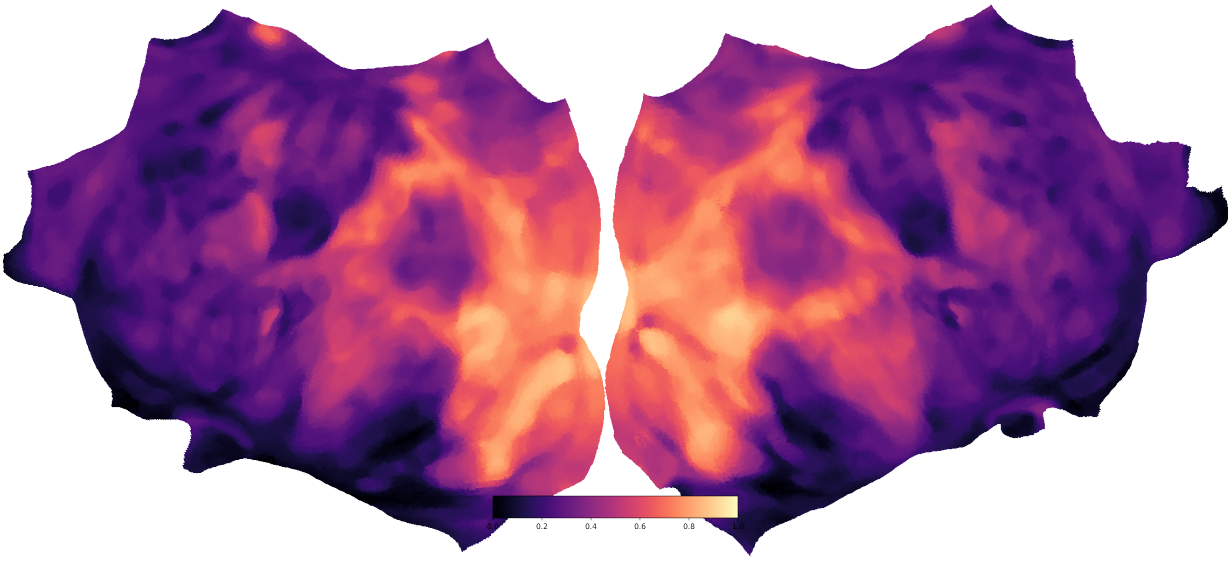


**Figure S27. Voxel-wise internal consistency (Methods B.1).**

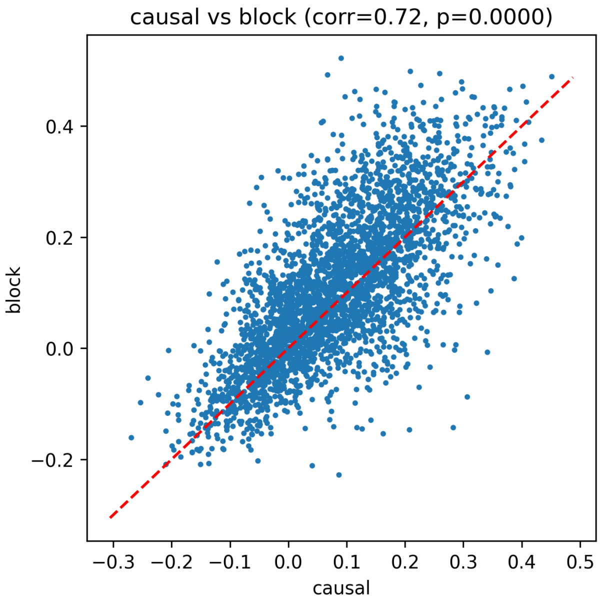


**Figure S28. Voxel-wise alignment score with causal inferencing vs. block inferencing.**

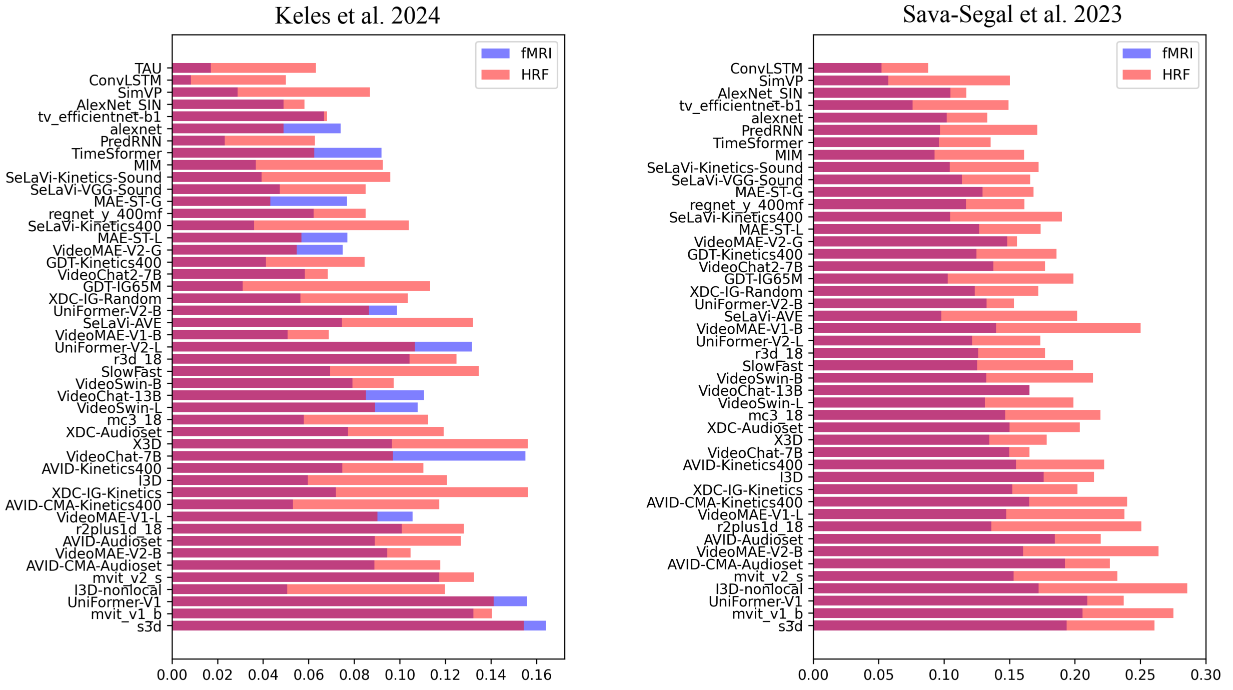


**Figure S29. FMRI alignment with / without the hemodynamic convolution:** the raw brain alignment of a subset of models on two datasets. The colours of transparent bars indicate two different activation processing methods: 1. Blue: model activations are simply delayed for 5 seconds to approximate the hemodynamic response delay; 2. Red: model activations are convolved with a hemodynamic function with 5 seconds as the peaking time parameter (see Methods). In general, convolving the hemodynamic function gives better alignment.
